# Centriolar satellites expedite mother centriole remodeling to promote ciliogenesis

**DOI:** 10.1101/2022.04.04.486992

**Authors:** Emma A. Hall, Dhivya Kumar, Suzanna L. Prosser, Patricia L. Yeyati, Vicente Herranz Pérez, Jose Manuel García Verdugo, Lorraine Rose, Lisa McKie, Daniel O. Dodd, Peter A. Tennant, Roly Megaw, Laura C. Murphy, Marissa Ferreira, Graeme Grimes, Lucy Williams, Tooba Quidwai, Laurence Pelletier, Jeremy F. Reiter, Pleasantine Mill

## Abstract

Centrosomes are orbited by centriolar satellites, dynamic multiprotein assemblies nucleated by PCM1. To study the requirement for centriolar satellites, we generated mice lacking PCM1. *Pcm1*^−/−^ mice display partially penetrant perinatal lethality with survivors exhibiting hydrocephalus, oligospermia and cerebellar hypoplasia, as well as variable expressivity of other ciliopathy features including cystic kidneys. *Pcm1*^−/−^ multiciliated ependymal cells and *PCM1*^−/−^ retinal pigmented epithelial 1 (RPE1) cells showed reduced ciliogenesis. *PCM1*^−/−^ RPE1 cells displayed reduced docking of the mother centriole to the ciliary vesicle and removal of CP110 and CEP97 from the distal mother centriole, indicating compromized early ciliogenesis. We show these molecular cascades are maintained *in vivo*, and we suggest that the cellular threshold to trigger ciliogenesis varies between cell types. We propose that PCM1 and centriolar satellites facilitate efficient trafficking of proteins to and from centrioles, inducing the departure of CP110 and CEP97 to initiate ciliogenesis.

## Introduction

A pair of microtubule-based centrioles form the heart of the centrosome. Centrosomes nucleate the mitotic spindle necessary for faithful chromosome segregation during cell division (***Nigg and Holland, 2018***). In most interphase cells, the older mother centriole matures into the basal body, which serves as the foundation for the single primary cilium, a specialized signaling antenna. In contrast, multiciliated cells lining the trachea and brain ependyma generate many basal bodies that nucleate the 10-100s of motile cilia per cell.

In all ciliated cells, dynamic remodeling of centrioles is required for ciliogenesis, the process of building a cilium. Key early steps consist of stepwise changes in the mother centriole, including acquisition of distal appendages and the removal from the distal end of the mother centriole of CP110 and CEP97, two proteins that inhibit assembly of the ciliary axoneme (***Čajánek and Nigg, 2014; Goetz et al., 2012; Schmidt et al., 2009, 2012; Sillibourne et al., 2013; Spektor et al., 2007; Tanos et al., 2013; Tsang et al., 2008***). How the cell remodels the mother centriole remains unclear.

Surrounding the centrosome and ciliary base are centriolar satellites, small membrane-less granules which move along cytoplasmic microtubules (***Bärenz et al., 2011; Kubo and Tsukita, 2003; Kubo et al., 1999; Odabasi et al., 2020***). Centriolar satellites assemble around a key component PCM1, necessary for centriolar satellite formation (***Dammermann and Merdes, 2002; Kubo and Tsukita, 2003; Odabasi et al., 2019; Wang et al., 2016***). A diverse array of proteins co-localize with PCM1 at centriolar satellites, many of which have been discovered through proteomic studies (***Gheiratmand et al., 2019; Gupta et al., 2015; Odabasi et al., 2019; Quarantotti et al., 2019***). Some centriolar satellite components also localize at centrioles themselves and include proteins critical to cilia formation and centriole duplication which, when mutated, cause human diseases such as ciliopathies and microcephaly (***Kodani et al., 2015; Lopes et al., 2011***). Centriolar satellites are dynamic, change in response to cell stresses, and have been implicated in diverse processes including Hedgehog signaling, autophagy, proteasome activity and aggresome formation (***Holdgaard et al., 2019; Hori and Toda, 2017; Joachim et al., 2017; Kubo and Tsukita, 2003; Lecland and Merdes, 2018; Odabasi et al., 2019; Prosser and Pelletier, 2020; Prosser et al., 2020; Tang et al., 2013; Villumsen et al., 2013; Wang et al., 2013a***). Recent work suggested that PCM1 is critical for maintaining neuronal cilia, disruption of which may cause schizophrenic-like features in aged mice (***Monroe et al., 2020***). Thus, understanding of the function of PCM1 and centriolar satellites is evolving.

To investigate the functions of centriolar satellites *in vivo*, we generated *Pcm1* null mice. We found that PCM1 is important for perinatal survival. *Pcm1*^−/−^ mice surviving the perinatal period displayed dwarfism, male infertility, hydrocephaly, cerebellar hypoplasia and partially expressive ciliopathy phenotypes such as hydronephrosis, reflecting important roles for centriolar satellites in promoting primary and motile ciliogenesis. In assessing how centriolar satellites promote ciliogenesis, we found that cells lacking PCM1 display compromized docking of the mother centriole to the ciliary vesicle and attenuated removal of CP110 and CEP97. Thus, we propose that centriolar satellites support ciliogenesis by removing CP110 and CEP97 from the mother centriole.

## Results

### *Pcm1* null mice display perinatal lethality and ciliopathy-associated phenotypes

To investigate the in vivo requirement and function of centriolar satellites in mammalian development, we used CRISPR/Cas9 to create deletions in mouse *Pcm1*. Among the mutations generated, *Pcm1*^Δ5−14^ introduced a frameshift after the first amino acid leading to a premature stop and *Pcm1*^Δ796−800^ caused a frameshift and premature stop in exon 6 (**Figure 1- figure supplement 1A**). Immunoblotting with antibodies to two regions of PCM1, PCM1 immunofluorescence of mouse embryonic fibroblasts (MEFs) derived from Pcm1 mutant mice, and mass spectrometry-based proteome analysis indicated that both mutations prevented formation of detectable PCM1 protein (**Figure 1A, B, Figure 1- figure supplement 1B, C**). Mice homozygous for either *Pcm1* mutation exhibited indistinguishable phenotypes (**Figure 1- figure supplement 2**). Thus, we surmise that both mutations are likely to be null and henceforth we refer to both alleles as *Pcm1*^−/^.

**Figure 1.**
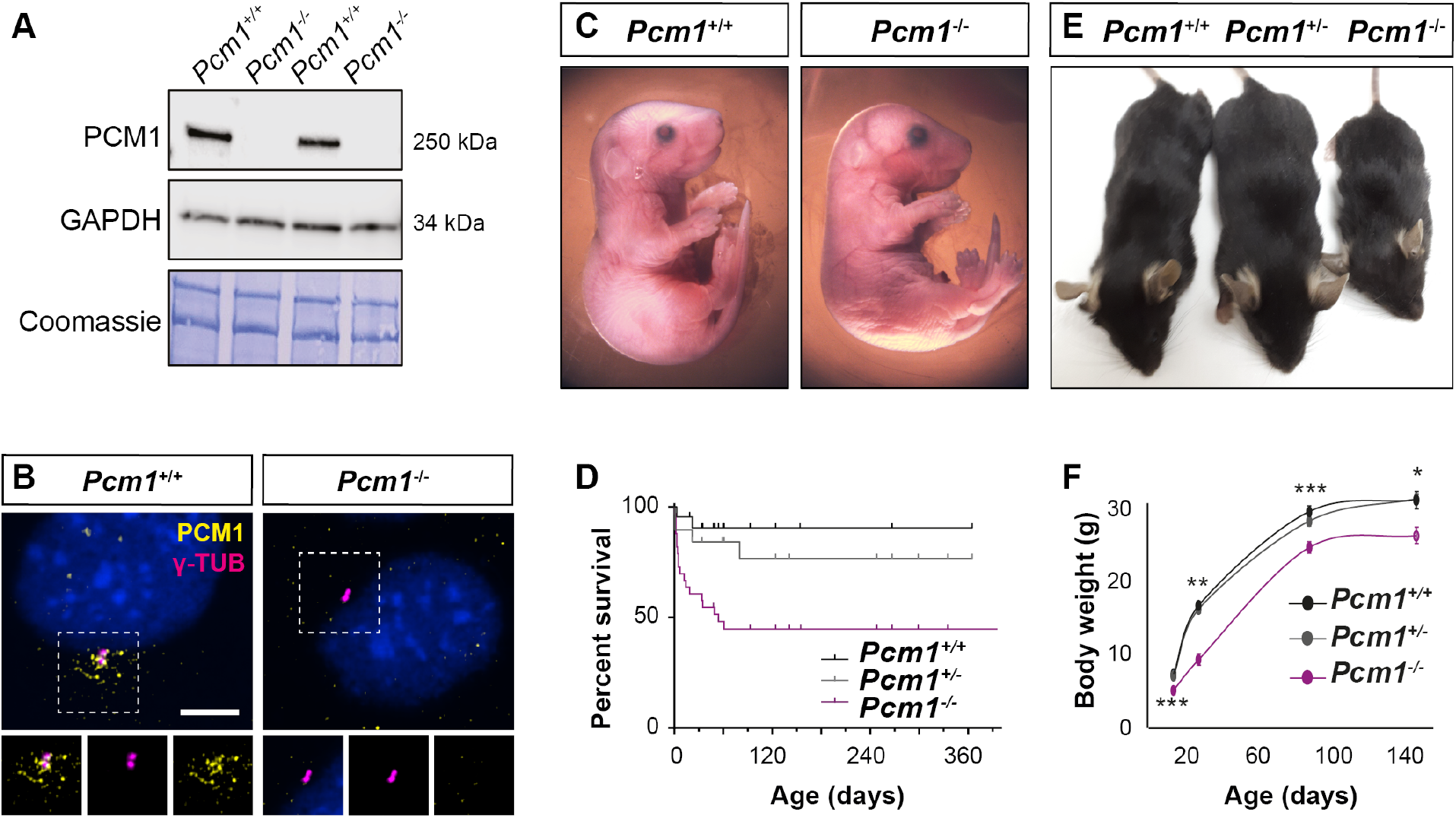
PCM1 is important for perinatal survival. **(A)** Immunoblot of MEF lysates from wild-type and *Pcm1*^−/−^ MEFs for PCM1 and GAPDH (loading control). Gel stained with Coomassie blue. *Pcm1*^−/−^ MEFs exhibit no detected PCM1. **(B)** Immunostaining reveals presence of PCM1 (yellow) in centriolar satellites around centrioles (*γ*tubulin, *γ*TUB, magenta) in wild-type MEFs but not in *Pcm1*^−/−^ MEFs. **(C)** E18.5 late gestation *Pcm1*^−/−^ neonates are grossly normal. **(D)** *Pcm1*^−/−^ mice display partially penetrant perinatal lethality. Kaplan-Meier graph showing that by P60, 50% of *Pcm1*^−/−^ mice have died either spontaneously or secondary to euthanasia for health concerns, mainly hydrocephaly. This wave of death occurs mostly between P0-P5, see **Figure 1 – figure supplement 1D. (E)** P28 surviving *Pcm1*^−/−^ mice are smaller than littermates. **(F)** Graph of mouse weights by age. *Pcm1*^−/−^ mice remain smaller than littermates. Student’s t-test * P<0.05, ** P<0.01, *** P<0.001. **Figure 1–Figure supplement 1. PCM1 promotes survival and growth**. **Figure 1–Figure supplement 2. *Pcm1***^Δ5−14^ **and *Pcm1***^Δ796−800^ **mice exhibit comparable phenotypes**.

*Pcm1*^−/−^ mice were present at normal Mendelian ratios at late gestation (embryonic day [E] 18.5) (**Figure 1C, D, Figure 1- figure supplement 1D**). As abrogation of cilia themselves results in midgestation lethality (***Huangfu et al., 2003***), this suggests that PCM1 is not essential for all ciliogenesis. However, by postnatal day (P) 5, most *Pcm1*^−/−^ mice died (**Figure 1D, Figure 1- figure supplement 1D**), revealing that PCM1 is important for perinatal survival.

Surviving *Pcm1*^−/−^ mice were smaller than littermate controls, weighing less than half of controls at P28 (**Figure 1E, F**). This dwarfism was detectable before birth, indicating intrauterine growth retardation (**Figure 1 - figure supplement 1E**). The brains of surviving *Pcm1*^−/−^ mice were proportionally smaller than those of littermates (**Figure 1- figure supplement 1F, G**), and displayed marked hydrocephaly (**Figure 2 A-C, Figure 2- figure supplement 1A, B**). Hydrocephaly can result from motile cilia dysfunction, suggesting that centriolar satellites may be required for cilia formation and/or function in ependymal cells.

**Figure 2.**
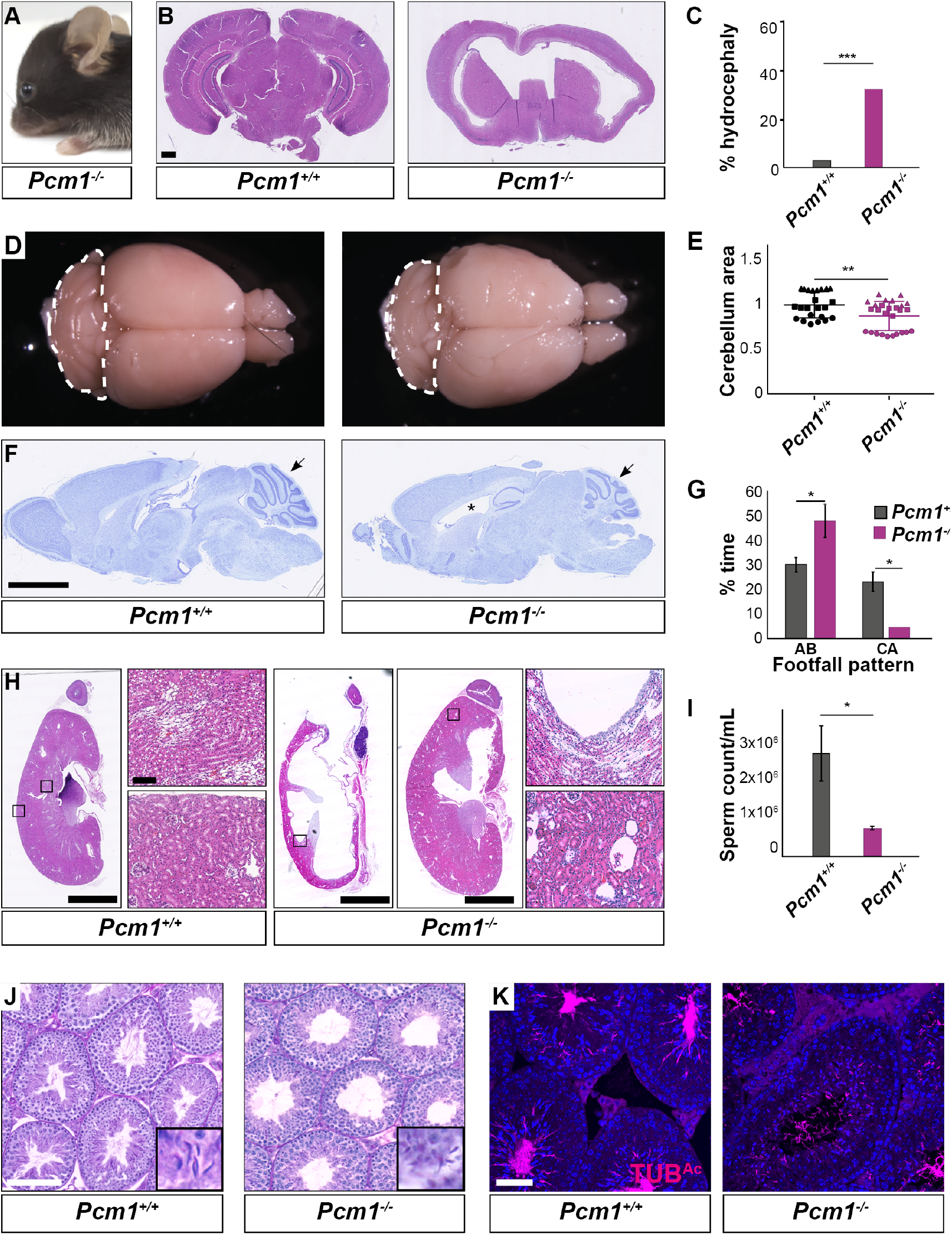
*Pcm1*^−/−^ mice display ciliopathy-associated phenotypes. **(A)** *Pcm1*^−/−^ mice display partially expressive hydrocephaly. **(B)** Coronal sections of 5-week-old wild type and *Pcm1*^−/−^ brains, revealing hydrocephaly. **(C)** Proportion of wild-type and *Pcm1*^−/−^ mice exhibiting hydrocephaly. **(D)** Gross morphology of 8-month-old wild-type and *Pcm1*^−/−^ brains reveals PCM1 promotes cerebellar growth. Cerebella are delineated with dotted lines. **(E)** Quantification of cerebellar area from sagittal wax sections of 2-8 month-old brains without frank hydrocephaly, normalized to the mean of wild type. n=3 each shape represents an animal. Mean +/-standard deviation. *P<0.05 Student’s t-test. **(F)** Cresyl violet-stained sagittal sections of 8-month-old brains. *Pcm1*^−/−^ brains are notable for cerebellar hypoplasia (arrows) and dilatated ventricles (*). **(G)** Catwalk gait analysis of adult wild-type and *Pcm1*^−/−^ mice. *Pcm1*^−/−^ mice show increased time in alternate (AB) gait and decreased time in cruciate (CA) gait. Mean +/-SEM, *Pcm1*^+/+^ n = 4. *Pcm1*^−/−^ n = 5. *P<0.05 Student’s t-test. **(H)** Kidney histology. A proportion of *Pcm1*^−/−^ mice display hydronephrosis (n=2/15). **(I)** Sperm count of wild-type and *Pcm1*^−/−^ mice. *Pcm1*^−/−^ mice display reduced sperm number. **(J)** Testis histology. Sperm are rarely observed in *Pcm1*^−/−^ tubules, indicating azoospermia. **(K)** Immunofluorescence staining of wild-type and *Pcm1*^−/−^ tubules for sperm flagella (acetylated tubulin, TUB^*Ac*^, magenta) and nuclei (DAPI, blue). *Pcm1*^−/−^ tubules contain few sperm. Scale bars represent 1 mm in B, 2.5 mm in F and H, 100 *μ*m in J, and 50 *μ*m in K. **Figure 2–Figure supplement 1. *Pcm1***^−/−^ **mice display a subset of ciliopathy-associated phenotypes**.

In the postnatal brain, primary cilia are critical for Hedgehog signaling in cerebellar granule cell precursors. Defects in cerebellar Hedgehog signaling attenuate expansion of the granule cell precursors (***Dahmane and Ruiz i Altaba, 1999; Spassky et al., 2008; Wallace, 1999; Wechsler-Reya and Scott, 1999***). The cerebella of *Pcm1*^−/−^ mice were smaller than those of littermate controls (**Figure 2D-F, Figure 2- figure supplement 1A**). As the cerebellum is important for motor coordination, we analyzed the gait of surviving *Pcm1*^−/−^ mice. Consistent with altered cerebellar function, *Pcm1*^−/−^ mice displayed ataxia (**Figure 2G**).

We investigated whether *Pcm1*^−/−^ mice exhibit other Hedgehog-associated phenotypes. A proportion of viable *Pcm1*^−/−^ mice (n=2/15) developed hydronephrosis (**Figure 2H**), which can also result from attenuated Hedgehog signaling (***Yu et al., 2002***).

Because retinal degeneration is characteristic of several ciliopathies and PCM1 was strongly expressed in the retina (**Figure 2- figure supplement 1 D**), we examined the retinas of *Pcm1*^−/−^ mice using fundal imaging and histological analysis at one year of age, as well as electroretinogram (ERG) at 9 months of age. *Pcm1*^−/−^ mice did not display characteristic features of photoreceptor death, such as changes to retinal pigmentation on fundoscopy or reduction of the outer nuclear layer on histology (**Figure 2- figure supplement 1 E, F**), or functional deficits (**Figure 2- figure supplement 1 G-I**). Therefore, PCM1 is not essential for photoreceptor survival, suggesting it is dispensable for photoreceptor ciliogenesis and ciliary trafficking.

Surviving *Pcm1*^−/−^ male mice were infertile with reduced sperm in seminiferous tubules (**Figure 2 I-K, Figure 2- figure supplement 2C**). The few *Pcm1*^−/−^ sperm identified exhibited disrupted head-to-tail coupling, abnormal manchettes and immobility (**Figure 2 - figure supplement 2C, Figure 2 – videos 1-3**). We previously discovered similar defects in male mice lacking centriolar satellite component CEP131 (also known as AZI1) (***Hall et al., 2013***), consistent with the idea that centriolar satellites are essential for mammalian spermatogenesis and male fertility. Thus, PCM1 promotes postnatal survival and is required for the function of multiple ciliated tissues.

### PCM1 promotes ciliogenesis in multiciliated cells

Ependymal cells lining the brain ventricles generate multiple motile cilia perinatally. Shortly after birth (P1), immature ependymal cells possess non-polarized, short cilia. Beginning at P3, ependymal cells form long, polarized cilia; this ciliogenesis occurs in a wave across the ventricle from caudal to rostral. By P15, ependymal cilia mature to generate metachronal rhythm (***Spassky et al., 2008***). Recent work showed that knockdown of *Pcm1* in cultured ependymal cells led to disrupted cilia ultrastructure and motility (***Zhao et al., 2021***).

To test whether defects in ependymal cilia could be the cause of hydrocephaly in *Pcm1*^−/−^ mice, we imaged ependymal cilia in lateral ventricle walls. By immunofluorescence, *Pcm1*^−/−^ mice exhibited numerous ependymal cell abnormalities, including fewer ependymal cells with multiple basal bodies at P3 and P5 (**Figure 3A-C, Figure 3- figure supplement 1A, B**). *Pcm1*^−/−^ mice also exhibited increased numbers of cells with rosette-like arrangements of basal bodies, representing a distinctive developmental stage in centriole biogenesis (**Figure 3B, F**). These results are consistent with a delay in centriole biogenesis in the absence of PCM1. In addition, basal bodies of *Pcm1*^−/−^ ependymal cells displayed disrupted orientation of basal feet and disrupted positioning of basal bodies within the apical domain (**Figure 3A, E, G, Figure 3- figure supplement 1D**). Basal foot misorientation can be caused by defective ciliary motility (***Guirao et al., 2010; Mirzadeh et al., 2010a***), whereas basal body positioning within the apical domain is thought to be independent ciliary motility, suggesting roles for PCM1 independent of motility (***Kishimoto and Sawamoto, 2012; Mirzadeh et al., 2010b***).

**Figure 3.**
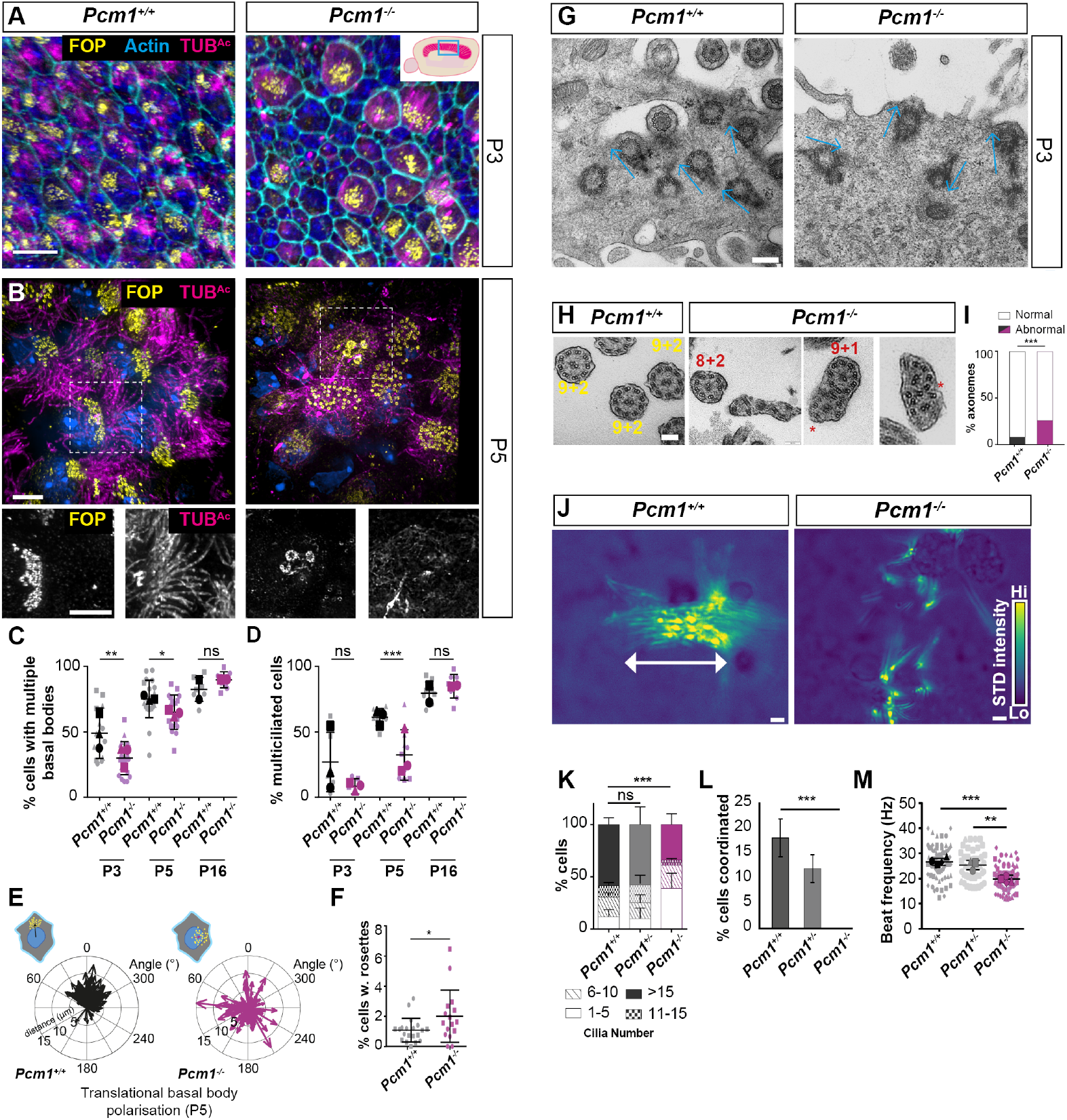
PCM1 is required for efficient multiciliogenesis. **(A)** Wild type and *Pcm1*^−/−^ P3 ventricle wholemounts immunostained for basal bodies (FOP, yellow), actin (phalloidin, cyan) and cilia (TUB^*Ac*^, magenta). Inset depicts area of ventricle imaged (cyan box). Basal body biogenesis and ciliogenesis are delayed in *Pcm1*^−/−^ ependymal cells. **(B)** P5 wild-type and *Pcm1*^−/−^ ventricle wholemounts immunostained for basal bodies (FOP, yellow), cilia (TUB^*Ac*^, magenta), and nuclei (DAPI, blue). Below: single optical planes highlight the persistence of deuterosome-like structures and the disruption of basal body planar polarization and ciliogenesis in *Pcm1*^−/−^ ependymal cells. **(C)** Quantification of number of P3, P5 and P16 wild-type and *Pcm1*^−/−^ ependymal cells with >4 basal bodies. Each shape represents an animal; the smaller symbols represent individual images and the larger shape the mean for each animal. Student’s t-test: * P<0.05, **: P<0.01, ns, not significant. Fewer *Pcm1*^−/−^ ependymal cells with multiple basal bodies are observed at P3 and P5, but not at P16. **(D)** Quantification of number of P3, P5 and P16 wild-type and *Pcm1*^−/−^ ependymal cells with multiple cilia. Each shape represents an animal; the smaller symbols represent individual images and the larger shape the mean for each animal. Student’s t-test: *** P<0.001, ns, not significant. **(E)** Rose plot of translational polarity of basal bodies in P5 wild-type and *Pcm1*^−/−^ ependymal wholemounts, as assessed from immunofluorescent images as in (A,B). The angle from the centre of the nucleus to the centre of the basal bodies is plotted relative to the average angle for that field of view, set at 0°C. **(F)** Quantification of P3 wild-type and *Pcm1*^−/−^ ependymal cells with deuterosome-like centriolar structures. Student’s t-test *P<0.05. **(G)** TEM of ependymal cells from P3 wild-type and *Pcm1*^−/−^ ventricles. Blue arrows indicate the direction from basal feet to centrioles. Basal feet are similarly oriented in wild-type cells but not in *Pcm1*^−/−^ cells. **(H)** TEM of ependymal cell cilia from P3 wild-type and *Pcm1*^−/−^ ventricles. Wild-type cilia display 9+2 microtubule arrangement. *Pcm1*^−/−^ cilia display axonemal defects, including missing microtubule doublets and axoneme fusion. **(I)** Quantification of P3 wild-type and *Pcm1*^−/−^ ependymal cilia structure. Chi squared test, *** P<0.001. **(J)** Colourized heat map (scale: yellow-high, blue-low) of maximum projection of the standard deviation of pixel intensity in **Figure 3 Video 1** and **2**, depicting wild-type and *Pcm1*^−/−^cultured ependymal cell cilia beating. Areas of high pixel intensity variation reflect areas of increased movement. Wild type ependymal cells display coordinated, planar beat pattern, that is absent in *Pcm1*^−/−^ ependymal cells. **(K)** Quantification of cilia number in cultured wild-type and *Pcm1*^−/−^ ependymal cells. Chi squared test, ***P<0.001, ns: not significant. *Pcm1*^−/−^ ependymal cells form fewer cilia. **(L)** Quantification of percentage of cultured wild-type and *Pcm1*^−/−^ ependymal cells with coordinated ciliary beating. Chi squared test, *** P<0.01. *Pcm1*^−/−^ependymal cells exhibit uncoordinated beating. **(M)** Cilia beat frequency of cultured wild-type and *Pcm1*^−/−^ ependymal cells. *Pcm1*^−/−^ cilia beat more slowly. Student’s t-test, ***P<0.001, **P<0.01. Scale bars: 15 *μ*m (A), 5 *μ*m (B), 200nm (G), 100nm (H) and 2 *μ*m (J). **Figure 3–Figure supplement 1. PCM1 is required for ciliary pocket formation and basal body polarization. Figure 3–Figure supplement 2. *Pcm1***^−/−^ **ependymal cells form elongated centriole-like structures**. **Figure 3–Figure supplement 3. Delayed expression of ciliary proteins in *Pcm1***^−/−^ **mTECs**.

Interestingly, *Pcm1*^−/−^ ependymal cells contained extremely elongated FOP-and Centrin-containing centriole-like structures measuring 5.0 ± 1.9 *μ*m (mean ± SD) in length (**Figure 3- figure supplement 2A-F**). At P3 *Pcm1*^−/−^ ependymal cell axonemes displayed ultrastructural defects, including missing microtubule doublets and fused axonemes (**Figure 3H and I**) with diminished ciliary pockets (**Figure 3- figure supplement 1C**).

To further analyze the function of PCM1 in multiciliogenesis, we cultured primary ependymal cells (***Guirao et al., 2010***) isolated from P0-P3 wild-type control and *Pcm1*^−/−^ mice. In culture, *Pcm1*^−/−^ ependymal cells formed fewer cilia than control ependymal cells (**Figure 3J, K**). High speed video microscopy revealed that *Pcm1*^−/−^ ependymal cilia beat slowly and uncoordinatedly (**Figure 3J, L, M, Figure 3 – videos 1-3**).

By P16, *Pcm1*^−/−^ mice possessed normal numbers of ependymal cells that were comparable to those of control littermates (**Figure 3C, D, Figure 3-supplement 1B**). Thus, PCM1 is not essential for ciliogenesis, but is required for timely basal body maturation and ciliogenesis in ependymal cells. Hydrocephaly in *Pcm1*^−/−^ mice is associated with delayed ependymal cell ciliogenesis and compromized ciliary motility.

Like the brain ventricles, the trachea is lined by motile multiciliated cells. To examine whether PCM1 also promotes ciliogenesis and ciliary motility in the airways, we examined mouse tracheal basal bodies and cilia by immunofluorescence. At P5, *Pcm1*^−/−^ tracheal multiciliated cells in vivo did not display grossly decreased numbers of cilia or basal bodies (**Figure 3- figure supplement 3B**).

To investigate the dynamics of ciliogenesis in these cells, we differentiated mouse tracheal epithelial cells (mTECS) into multiciliated cells in vitro (***Eenjes et al., 2018; You et al., 2002***). Concurring with a previous reports on the lack of effect of *Pcm1* depletion in mTECs (***Vladar and Stearns, 2007***) *Pcm1*^−/−^ mTECs displayed no clear alteration in basal body biogenesis or ciliogenesis (**Figure 3- figure supplement 3A**). However, proteomic analysis of mTEC maturation revealed that many motile ciliary proteins, including dynein motors, dynein assembly factors and dynein docking factors, were reduced at early ciliogenenic stages (ALI day 7) in *Pcm1*^−/−^ mTECs (**Figure 3- figure supplement 3C**). Similar to the recovery of cell morphogenesis we observed in *Pcm1*^−/−^ ependymal cells, proteomic differences in *Pcm1*^−/−^ mTECs resolved by ALI day 21 (**Figure 3- figure supplement 3C**). Thus, as in ependymal cells, PCM1 promotes timely ciliogenesis in tracheal cells.

### PCM1 is required for centriolar satellite integrity

To assess whether PCM1 is essential for centriolar satellite integrity, we analyzed *Pcm1*^−/−^ MEFs and *PCM1*^−/−^ RPE1 cells (***Kumar et al., 2021***). Immunoblot and immunofluorescence analyses confirmed loss of PCM1 protein in the mutant cells (**Figure 1A, B, Figure 4G, H**). In addition to PCM1, centriolar satellites contain proteins such as the E3 ligase MIB1 and CEP131 (***Hall et al., 2013; Staples et al., 2012; Villumsen et al., 2013***). In control MEFs, CEP131 and MIB1 localized to both centriolar satellites and to the centrioles themselves. In *Pcm1*^−/−^ MEFs, centriolar satellite CEP131 and MIB1 were absent and CEP131 displayed increased accumulation at centrioles (**Figure 4A-D**). Similarly, in control RPE1 cells, CEP131 and CEP290 localized to both centriolar satellites and to the centrioles themselves. In *PCM1*^−/−^ RPE1 cells, centriolar satellite proteins CEP131 and CEP290 were absent and both displayed increased accumulation at centrioles (**Figure 4E, F**). We conclude that PCM1 is critical for centriolar satellite integrity. In the absence of satellites, some satellite proteins (e.g., CEP131, CEP290) over-accumulate at centrioles, while others (e.g., MIB1) do not, highlighting the protein-specific role centriolar satellites play in controlling centriolar localization. We propose that centriolar satellites both deliver and remove select cargos from centrioles.

**Figure 4.**
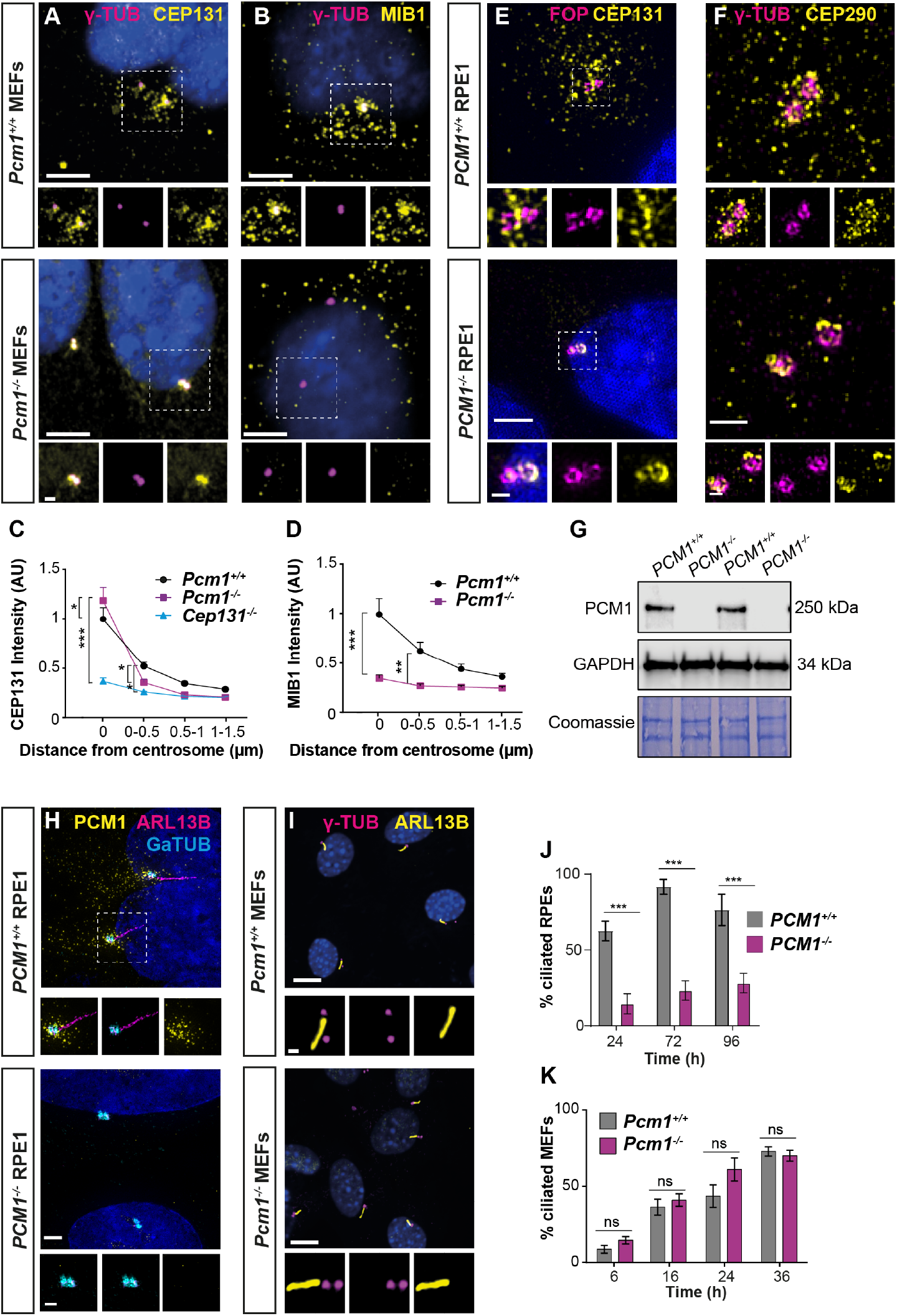
PCM1 is essential for centriolar satellite integrity and, in some cells, ciliogenesis. **(A)** Wild-type and *Pcm1*^−/−^ MEFs immunostained for CEP131 (yellow), centrioles (*γ*-tubulin, magenta), and nuclei (DAPI, blue). **(B)** As in (A), except MIB1 immunostaining is in yellow. **(C)** Quantification of CEP131 intensity in concentric rings round the centrosome. *Cep131*^−/−^ MEFs are included as a control ***Hall et al***. (***2013***). 2-way ANOVA, comparing WT to mutant, Dunnett correction for multiple testing * P < 0.05, ** P < 0.01, *** P < 0.001. Error bars represent standard error of the mean. In the absence of PCM1, CEP131 relocalizes from centriolar satellites to the centrosome. **(D)** Quantification of MIB1 intensity as in (C). Without PCM1, MIB1 disperses throughout the cell. **(E)** Wild-type and *PCM1*^−/−^ RPE1 cells immunostained for CEP131 (yellow), centrioles (FOP, magenta), and nuclei (DAPI, blue). In the absence of PCM1, CEP131 no longer localizes to the satellites and persists at centrioles. **(F)** Wild-type and *PCM1*^−/−^ RPE1 cells immunostained for CEP290 (yellow), centrioles (*γ*-tubulin, magenta), and nuclei (DAPI, blue). PCM1 is also required for CEP290 satellite localization and dispensable for its centriolar localization. **(G)** Immunoblot of wild-type and *PCM1*^−/−^ RPE1 cells for PCM1 and GAPDH (loading control). Gel stained with Coomassie blue. **(H)** Wild-type and *PCM1*^−/−^ RPE1 cells immunostained for PCM1 (yellow), cilia (ARL13B, magenta), centrioles (*γ*-tubulin, cyan) and nuclei (DAPI, blue). In *PCM1*^−/−^ RPE1 cells, PCM1 is undetectable and ciliogenesis is attenuated. **(I)** Wild-type and *Pcm1*^−/−^ MEFs immunostained for cilia (ARL13B, yellow), centrioles (*γ*-tubulin, magenta) and nuclei (DAPI, blue). PCM1 is not critical for ciliogenesis in MEFs. (J) Quantification of ciliogenesis in wild-type and *PCM1*^−/−^ RPE1 cells serum starved for 24, 72 and 96 h. Bar graphs show mean ± SD. Asterisks represent p < 0.001 determined using unpaired students t test. n>100 cells from 3 biological replicates. (K) Quantification of ciliogenesis in wild-type and *Pcm1*^−/−^ MEFs serum starved for 6-36 h. Students t-test, ns = not significant. Scale bars: 5 *μ*m (A, B), 1 *μ*m (A, B insets), 2 *μ*m (E), 1 *μ*m (F), 0.5*μ*m (E, F insets), 10 *μ*m (H, I) and 1 *μ*m (H, I insets). **Figure 4–Figure supplement 1. PCM1 is dispensable for ciliogenesis in MEFs**.

One way in which satellites could traffic cargos to and from centrioles would be via their movement within the cell. To visualize PCM1, we engineered mouse *Pcm1* to express a fusion of PCM1 and the SNAP tag from the endogenous locus. We derived MEFs from *Pcm1*^*SNAP*^ mice and covalently labeled PCM1-SNAP with tetramethylrhodamine (TMR) (***Crivat and Taraska, 2012***) and imaged centriolar satellite movement relative to cilia. Consistent with previous reports (***Conkar et al., 2019***), centriolar satellites moved both towards and away from the ciliary base, with frequent fission and fusion at the ciliary base (**Figure 4 – video 1**).

To further explore how centriolar satellites promote ciliogenesis, we examined ciliogenesis in MEFs and RPE1 cells lacking PCM1. In accordance with previous observations (***Odabasi et al., 2019; Wang et al., 2016***), ciliogenesis was abrogated in *PCM1*^−/−^ RPE1 cells examined 24 h post-serum deprivation (Figure 4H, J). The number of ciliated *PCM1*^−/−^ RPE1 cells increased slightly under prolonged serum deprivation, indicating that centriolar satellites promote timely ciliogenesis in RPE1 cells (**Figure 4J**). In marked contrast, and consistent with the tissue-specific effects on ciliogenesis in *Pcm1*^−/−^ embryos, ciliogenesis was not perturbed in *Pcm1*^−/−^ MEFs, with *Pcm1*^−/−^ MEFs displaying cilia number, centrosome number and cilia length indistinguishable from those of controls (**Figure 4I, K, Figure 4-Figure supplement 1 A-D**). Thus, PCM1 plays cell-type specific roles in ciliogenesis, despite broad roles in adjusting the centriolar localization of proteins such as CEP131, leading us to investigate how PCM1 and centriolar satellites promote ciliogenesis in certain cell types.

### PCM1 is dispensable for distal and sub-distal appendage assembly and removal of Centrobin

Among the multiple early steps of ciliogenesis are the removal of daughter centriole-specific proteins such as Centrobin and Talpid3, which localize to the distal centriole and are required for ciliogenesis (***Stephen et al., 2015; Wang et al., 2018***). A previous study proposed a role for PCM1-positive centriolar satellites in regulating the abundance of Talpid3 (***Wang et al., 2016***). We found that Talpid3 localization in *PCM1*^−/−^ RPE1 and its abundance at *PCM1*^−/−^ centrioles were equivalent to those of controls (**Figure 5- figure supplement 1A, B**). Talpid3 is implicated in the removal of Centrobin from the mother centriole (***Wang et al., 2018***). In *PCM1*^−/−^ RPE1 cells, as in control cells, Centrobin localized specifically to the daughter centriole (**Figure 5- figure supplement 1C, D**). Thus, Talpid3 recruitment to centrioles and Centrobin removal from the mother centriole are not dependent upon PCM1 or, by extension, centriolar satellites.

Distal appendages anchor the mother centriole to the ciliary membrane and sub-distal appendages position the cilium within cells (***Mazo et al., 2016; Schmidt et al., 2012; Sillibourne et al., 2013; Tanos et al., 2013***). Since centriolar satellite cargos (e.g., CEP90, OFD1 and MNR) are essential for ciliogenesis and distal appendage assembly (***Kumar et al., 2021***), we hypothesized that PCM1 may participate in distal or sub-distal appendage formation. To test this hypothesis, we examined localization of components of the distal (i.e., FBF1 and ANKRD26) and sub-distal appendages (i.e., Ninein) at the mother centriole. In *PCM1*^−/−^ RPE1 cells, both distal and sub-distal appendage components localized to the mother centriole (**Figure 5- figure supplement 1 E-J**), although we observed a small but significant decrease in the levels of distal appendage proteins at the mother centriole. Serial section transmission electron microscopy (TEM) confirmed that sub-distal and distal appendages were present in *PCM1*^−/−^ RPE1 cells (**Figure 5- figure supplement 2**). Therefore, centriolar satellites are not required for the assembly of distal or sub-distal appendages at the mother centriole.

### PCM1 promotes formation of the ciliary vesicle

After acquiring distal appendages, the mother centriole docks to preciliary vesicles (PCVs), small vesicles which accumulate at the distal appendages of the mother centriole and are converted into a larger ciliary vesicle (CV) (***Schmidt et al., 2012; Sillibourne et al., 2013; Tanos et al., 2013***). To further examine the cause of reduced ciliogenesis in RPE1 cells lacking centriolar satellites, we investigated whether PCV docking or ciliary vesicle formation depends on PCM1.

Myosin-Va marks preciliary and ciliary vesicles (***Wu et al., 2018***). Using 3D-SIM imaging of Myosin-Va, we identified ciliary vesicles at the basal bodies of control RPE1 cells 1 h after we had induced ciliogenesis. In contrast, *PCM1*^−/−^ RPE1 cells showed reduced Myosin-Va at ciliary vesicles (**Figure 5A, B**), suggesting that centriolar satellites promote timely centriolar docking of PCVs.

**Figure 5.**
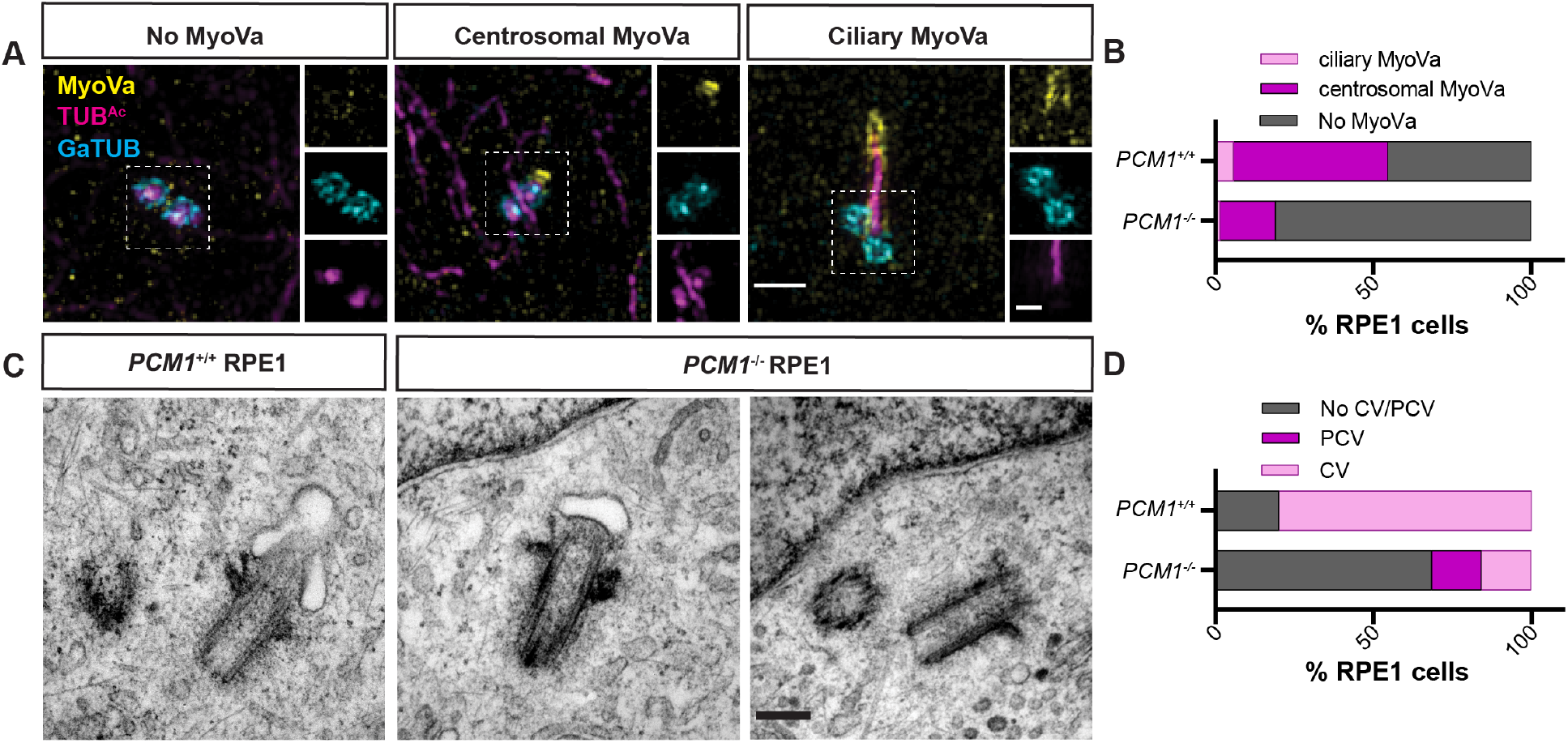
PCM1 promotes mother centriole docking to preciliary vesicles. **(A)** 3D-SIM imaging of Myosin-Va (MyoVa, yellow), centrioles (*γ*-tubulin, cyan) and cilia (TUB^*Ac*^, magenta) in wild-type and *PCM1*^−/−^ RPE cells 1 h post serum starvation. Scale bar is 1 *μ*m and 0.5 *μ*m for main panels and insets, respectively. **(B)** Quantification of MyoVa staining pattern in WT and *PCM1*^−/−^ RPE1 cells as exemplified in A. **(C)** Serial-section TEM of RPE1 cells in early ciliogenesis (serum starved for 1 h). Scale bar is 200 nm. **(D)** Quantification of basal bodies associated with ciliary vesicles (CV) and preciliary vesicles (PCV) in WT and *PCM1*^−/−^ RPE1 cells from TEM images. Ciliary and preciliary vesicle recruitment to the mother centriole is reduced in the absence of PCM1. **Figure 5–Figure supplement 1. PCM1 is dispensable for mother centriole maturation. Figure 5–Figure supplement 2. PCM1 promotes mother centriole association with vesicles**.

To assess centriolar docking to the ciliary vesicle formation using a complementary approach, we performed serial section TEM of control and *PCM1*^−/−^ RPE1 cells in which we had induced ciliogenesis by serum starvation for 1 h. We quantified PCVs and CVs at mother centrioles. In *PCM1*^−/−^ cells, mother centrioles (identified by the presence of distal and sub-distal appendages) exhibited reduced association with both PCVs and CVs (**Figure 5C, D, Figure 5- figure supplement 2**). Thus, TEM confirms that centriolar satellites promote the attachment of the mother centriole to preciliary vesicles, a critical early step in ciliogenesis.

### PCM1 promotes CP110 and CEP97 removal from the mother centriole

In vertebrates, CP110 is required for docking of the mother centriole to preciliary vesicles (***Wa-lentek et al., 2016; Yadav et al., 2016***) and is removed from the mother centriole subsequent to docking (***Lu et al., 2015; Wu et al., 2018***). The CP110 and CEP97 cap inhibits ciliogenesis, and their removal from the distal mother centriole is important for axoneme elongation (***Spektor et al., 2007; Yadav et al., 2016***). Since PCM1 promotes timely ciliary vesicle formation, we examined whether CP110 and CEP97 removal also requires PCM1. In contrast to control cells, CP110 and CEP97 persisted at the distal mother centriole in *PCM1*^−/−^ RPE1 cells (**Figure 6A, B, E, F**).

**Figure 6.**
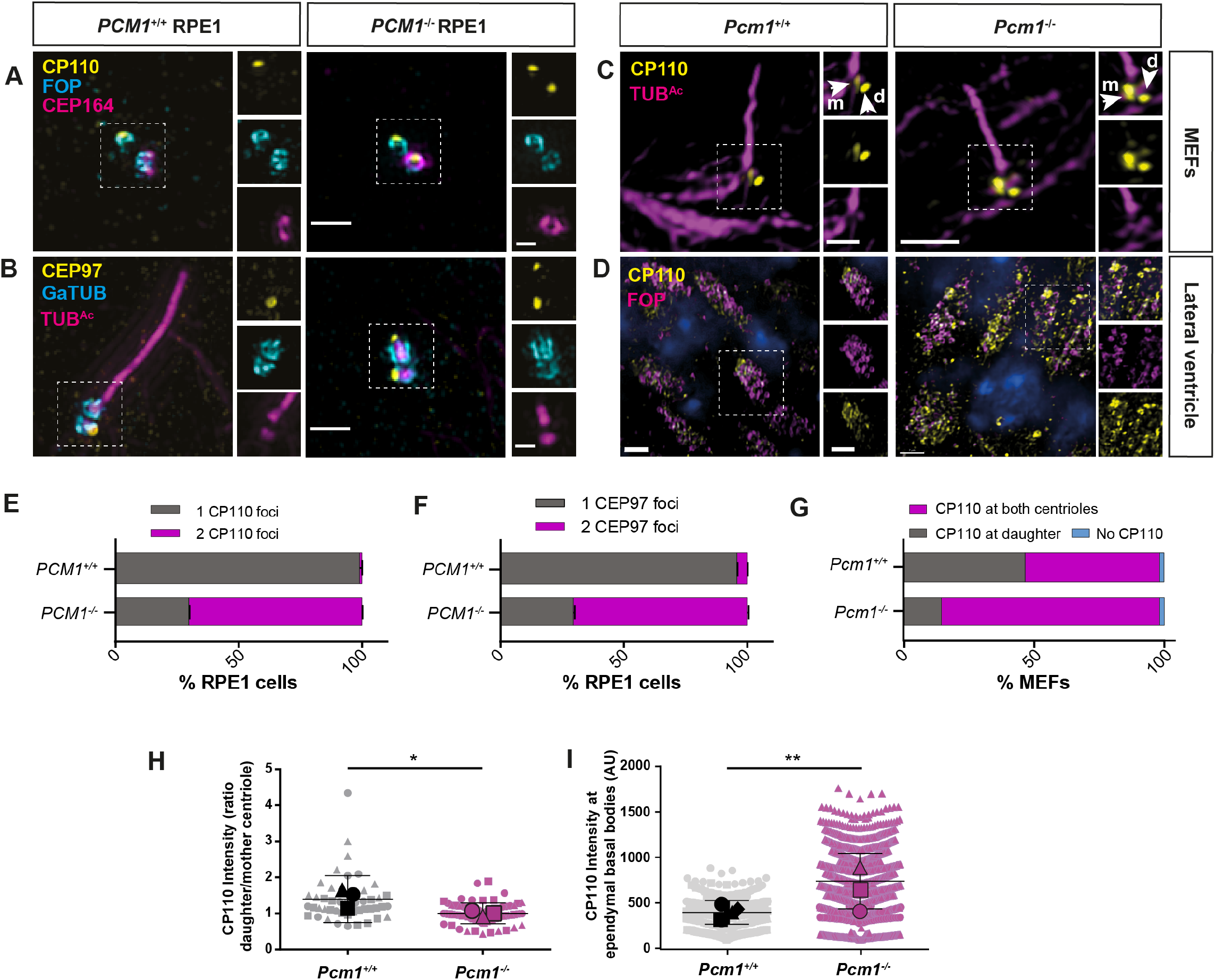
PCM1 promotes mother centriole docking to preciliary vesicles. **(A)** Wild-type and *PCM1*^−/−^ RPE1 cells serum starved for 24h immunostained for CP110 (yellow), centrioles (FOP, cyan), and distal appendages (CEP164, magenta). PCM1 is required to remove CP110 from the mother centriole (identified by distal appendages) after 24 h serum starvation. **(B)** Wild-type and *PCM1*^−/−^ RPE1 cells serum starved for 24h immunostained for CEP97 (yellow), centrioles (*γ*-tubulin, cyan), and cilia (TUB^*Ac*^, magenta). **(C)** Wild-type and *Pcm1*^−/−^ MEFs serum starved for 24 h immunostained for CP110 (yellow) and cilia (TUB^*Ac*^, magenta). After 24 h serum starvation, some CP110 remains on the mother (m) centriole in wild type MEFs (identified by cilium) and is lower than on the daughter (d) centriole. PCM1 is required to reduce CP110 levels at the mother centriole. **(D)** Wild-type and *Pcm1*^−/−^lateral ventricular wall immunostained for CP110 (yellow), centrioles (FOP, cyan), and nuclei (DAPI, blue). PCM1 is required to reduce CP110 at ependymal cell basal bodies. **(E)** Percentage of RPE1 cells with CP110 levels at one or two centrioles. **(F)** Percentage of RPE1 cells with CEP97 levels at one or two centrioles. **(G)** Percentage of MEFs serum starved for 24 h with CP110 levels at none, one or two centrioles. **(H)** The ratio of CP110 intensity on daughter and mother centrioles in wild-type and *Pcm1*^−/−^MEFs serum starved for 24 h. The lower ratio in *Pcm1*^−/−^ MEFs suggests that PCM1 reduces CP110 at the mother centriole. **(I)** Intensity of CP110 intensity in wild-type and *Pcm1*^−/−^ ependymal cells. Student’s t-test, * P<0.05, ** P < 0.01, *** P<0.001. Scale bars in A, B represent 1 *μ*m and 0.5 *μ*m in main panels and insets, in C represent 5 *μ*m and 1 *μ*m in main panels and insets, and in D represent 2 *μ*m.

In wild-type MEFs, a small amount of CP110 persisted on the mother centriole even after axoneme elongation (**Figure 6 C, G, H**). Interestingly, despite undergoing ciliogenesis at rates equal to that of wild-type cells, mother centrioles in *Pcm1*^−/−^ MEFs had CP110 levels comparable to daughter centrioles after 24 h serum starvation (**Figure 6C, G, H**). Thus, PCM1 is essential for removing CP110 from the mother centriole, but CP110 removal is not required for ciliogenesis in MEFs.

In *Pcm1*^−/−^ ependymal cells *in vivo*, CP110 levels were elevated at P3, an age when ependymal calls are engaged in ciliogenesis (**Figure 6D, I**). Thus, in diverse cell types, some of which depend on PCM1 to support ciliogenesis and some of which do not, PCM1 is required to remove CP110 from the mother centriole.

### PCM1 promotes transition zone formation and IFT recruitment

Following ciliary vesicle docking and removal of CP110 and CEP97 from the mother centriole, ciliogenesis proceeds by recruiting IFT machinery and building the transition zone (***Ishikawa and Marshall, 2011***). Since PCM1 promotes ciliary vesicle docking and CP110 and CEP97 removal, we hypothesized that subsequent recruitment of IFT and transition zone components would be compromized in cells lacking PCM1.

To test this hypothesis, we immunostained control and *PCM1*^−/−^ RPE1 cells with antibodies to IFT88 and IFT81. As expected, IFT88 and IFT81 localized to mother centrioles and along the length of cilia in control cells (**Figure 7A, C**). Localization of both IFT88 and IFT81 at mother centrioles was reduced in *PCM1*^−/−^ RPE1 cells (**Figure 7A-D**), suggesting that IFT recruitment to the mother centriole is promoted by centriolar satellites. In contrast, ciliary and basal body levels of IFT88 were normal in *Pcm1*^−/−^ MEFs (**Figure 7- figure supplement 1A, B**), suggesting that in cell types that ciliate normally without PCM1, IFT is recruited to mother centrioles independently of PCM1.

**Figure 7.**
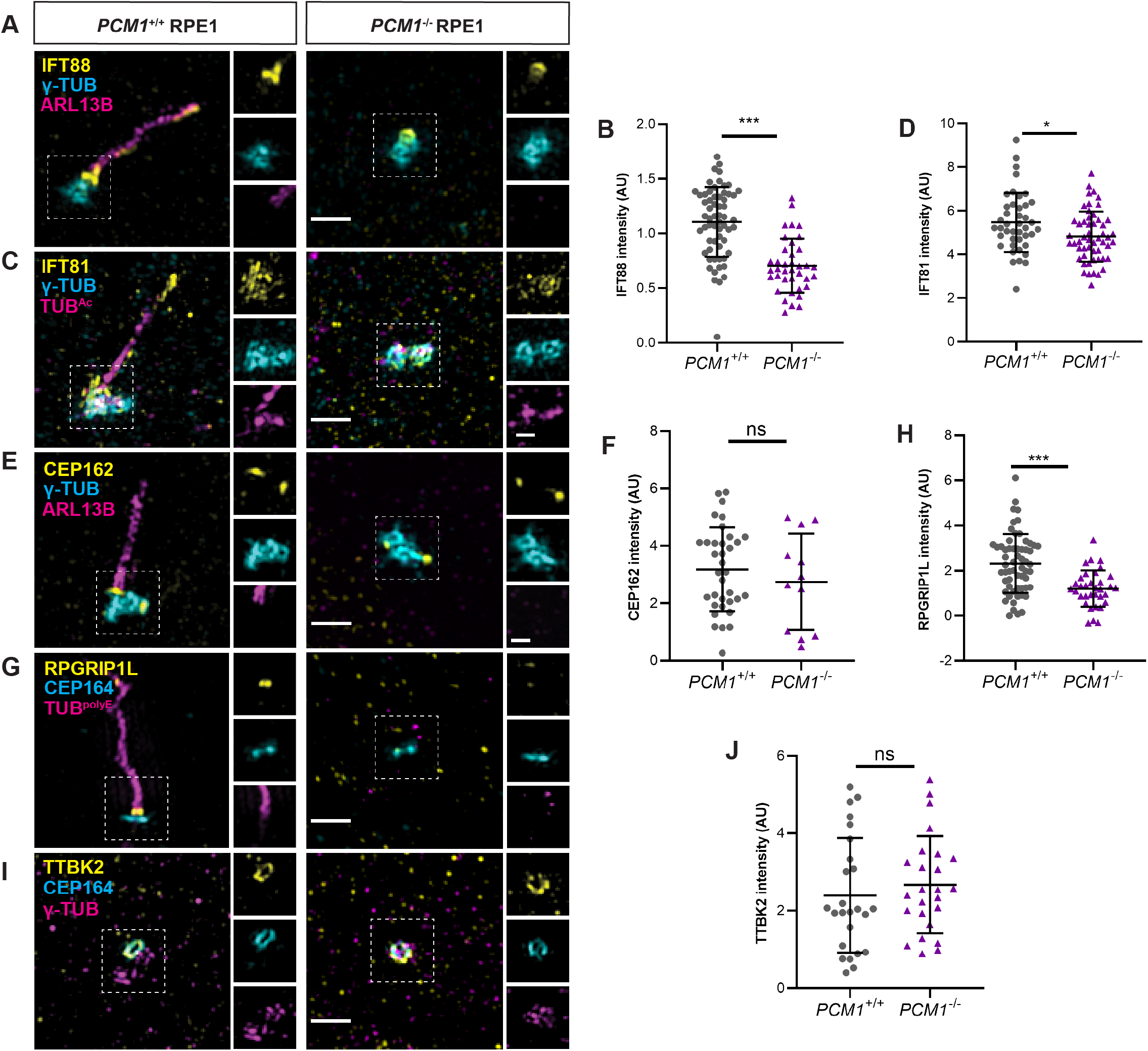
PCM1 promotes IFT recruitment and transition zone formation. **(A)** Wild-type and *PCM1*^−/−^ RPE1 cells immunostained for IFT88 (yellow), centrioles (*γ*-tubulin, cyan), and cilia (ARL13B, magenta). **(B)** Quantification of IFT88 intensity at basal bodies. **(C)** Immunostaining for IFT81 (yellow), centrioles (*γ*-tubulin, cyan), and cilia (TUB^*Ac*^, magenta). **(D)** Quantification of IFT88 intensity at basal bodies. IFT-B recruitment to basal bodies is reduced in *PCM1*^−/−^ RPE1 cells. **(E)** Immunostaining for CEP162 (yellow), centrioles (*γ*-tubulin, cyan), and cilia (ARL13B, magenta). **(F)** Quantification of CEP162 intensity at basal bodies. PCM1 is dispensable for recruiting CEP162 to basal bodies. **(G)** Immunostaining for RPGRIP1L (yellow), distal appendages (CEP164, cyan), and cilia (TUB^*P olyE*^, magenta). **(H)** Quantification of RPGRIP1L intensity at transition zones. RPGRIP1L recruitment to transition zones is reduced in *PCM1*^−/−^ RPE1 cells. **(I)** Immunostaining for TTBK2 (yellow), distal appendages (CEP164, cyan), and centrioles (*γ*-tubulin, magenta). **(J)** Quantification of TTBK2 intensity at basal bodies. PCM1 is dispensable for recruiting TTBK2 to basal bodies. Scale bars in main figures represent 1 *μ*m and in insets represent 0.5 *μ*m. Student’s t-test, * P<0.05, *** P<0.001, ns: not significant. **Figure 7–Figure supplement 1. PCM1 does not control IFT88 levels in MEF cilia**.

The transition zone controls ciliary protein composition. We determined whether PCM1 was required for the formation of the transition zone by assessing the localization of CEP162, an axoneme-associated protein that recruits components of the transition zone such as RPGRIP1L (***Wang et al., 2013b***). Recruitment of CEP162 to the mother centriole was unaffected in *PCM1*^−/−^ RPE1 cells (**Figure 7E, F**). In contrast, *PCM1*^−/−^ RPE1 cells exhibited reduced RPGRIP1L at the transition zone (**Figure 7G, H**). Therefore, centriolar satellites promote both IFT recruitment and transition zone formation at the RPE1 cell mother centriole.

### Centriolar satellites restrict CP110 and CEP97 levels at centrioles

To explore the mechanisms by which centriolar satellites regulate CP110 and CEP97 levels at the centrioles, we examined the localization of TTBK2. TTBK2 is a kinase recruited by CEP164, a distal appendage component required to remove CP110 and CEP97 from mother centrioles (***Goetz et al., 2012***). In *PCM1*^−/−^ RPE1 cells, TTBK2 recruitment to distal mother centrioles was equivalent to that of control cells (**Figure 7I, J**). These results suggest that centriolar satellites regulate CP110 and CEP97 removal from the distal mother centriole through a mechanism independent of TTBK2 recruitment.

As PCM1 is dispensable for the localization of TTBK2 at the distal mother centriole, we considered alternative mechanisms by which PCM1 may regulate local CP110 and CEP97 levels at the mother centriole. Since centriolar satellites are highly dynamic and localization of CP110 and CEP97 is actively controlled at the initiation of ciliogenesis, we hypothesized that CP110 and CEP97 are transported away from the centrioles via satellites. A prediction of this model is that CP110 and CEP97 should localize to satellites.

We examined RPE1 cells for CP110 and CEP97 and found that, indeed, CP110 and CEP97 colocalized with PCM1 and CEP290 at centriolar satellites in cycling cells (**Figure 8A, B, Figure 8- figure supplement 1A, B**). Moreover, this satellite pool of CP110 was absent in *PCM1*^−/−^ RPE cells (**Figure 8- figure supplement 1B**). Consistent with CP110 and CEP97 co-localizing with PCM1 at centriolar satellites, CP110 and CEP97 co-immunoprecipitated with PCM1 in cycling cells (**Fig 8I**).

**Figure 8.**
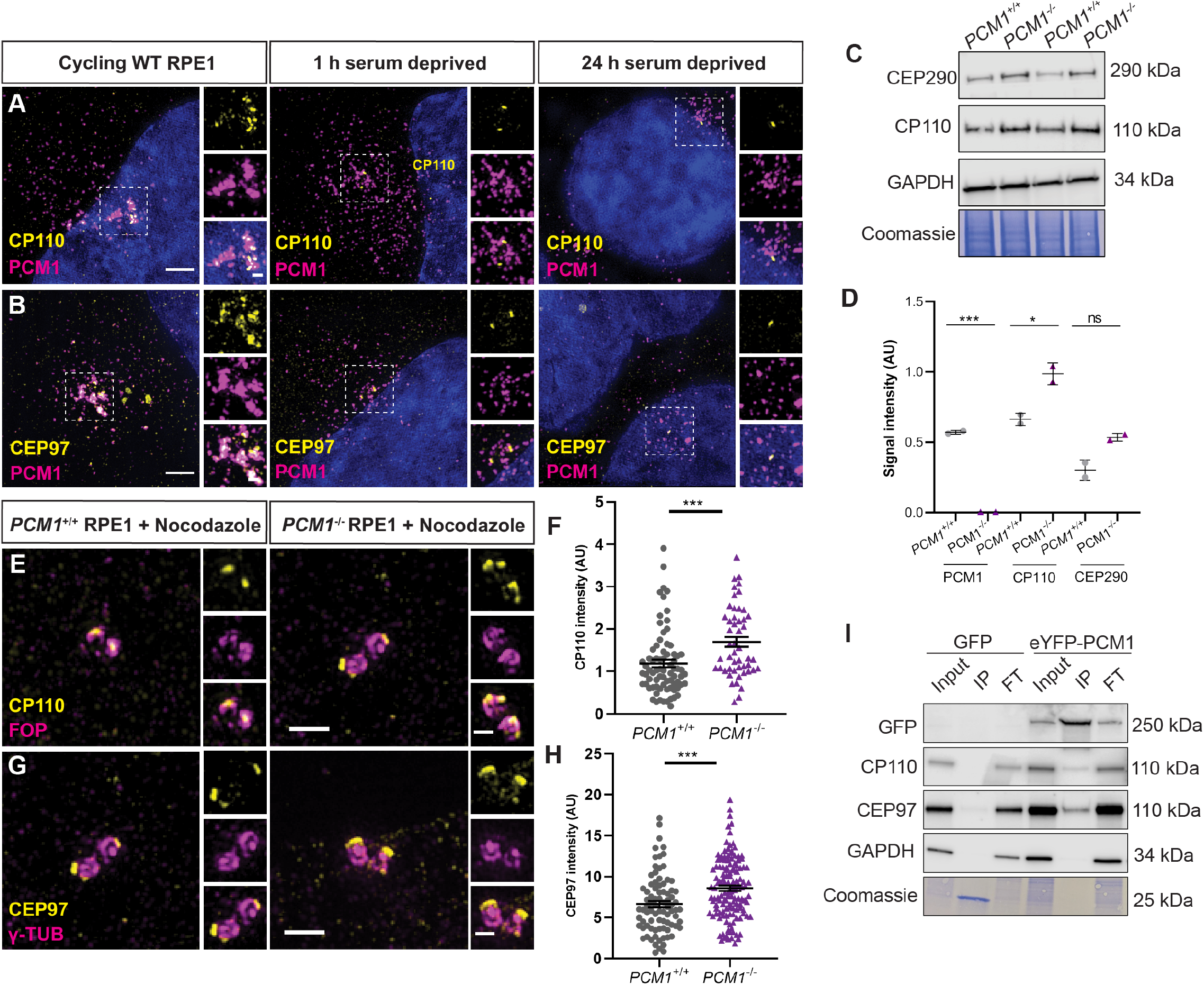
PCM1 restricts CP110 and CEP97 localization to distal mother centrioles. **(A)** Wild-type and *PCM1*^−/−^ RPE1 cells immunostained for CP110 (yellow), centriolar satellites (PCM1, magenta), and nuclei (DAPI, blue) in cells with serum, 1 h or 24 h after withdrawing serum. CP110 localizes to satellites in cycling cells, and leaves satellites within an hour of initiating ciliogenesis. **(B)** Immunostaining for CEP97 (yellow), centriolar satellites (PCM1, magenta), and nuclei (DAPI, blue). CEP97 also localizes to satellites in cycling cells and leaves satellites within an hour of initiating ciliogenesis. **(C)** Immunoblot of wild-type and *PCM1*^−/−^ RPE1 cell lines lysates for CP110 and GAPDH, as well as Coomassie stain of gels. Cells were deprived of serum for 24 h prior to lysis. **(D)** Quantification of PCM1 and CP110 levels from immunoblots. In the absence of PCM1, CP110 levels are increased. **(E)** Wild-type and *PCM1*^−/−^ RPE1 cells immunostained for CP110 (yellow) and centrioles (FOP, magenta). Cells were treated with nocodazole to disperse the centriolar satellite pool of CP110, leaving the centriolar pool. **(F)** Quantification of CP110 levels at centrioles stained as in E. PCM1 restricts CP110 localization to centrioles. **(G)** Immunostaining for CEP97 (yellow) and centrioles (*γ*-tubulin, magenta). **(H)** Quantification of CEP97 levels at centrioles stained as in G. PCM1 restricts CEP97 localization to centrioles. **(I)** Total cell lysates of *PCM1*^−/−^ RPE1 cell lines stably expressing eGFP or eYFP-PCM1 subjected to immunoprecipitation with anti-GFP. Precipitating proteins were immunoblotted for GFP, CP110, CEP97 and GAPDH. Coimmunoprecipitation reveals that eYFP-PCM1 interacts with CP110 and CEP97. IP is eluate and FT is flow through. Scale bars represent 1 *μ*m and 0.5 *μ*m in main panels and insets respectively. Student’s t-test * P<0.05, *** P<0.001, ns: not significant. **Figure 8–Figure supplement 1. CP110 localizes to satellites in a CEP290-dependent manner**.

By examining RPE1 cells at different timepoints after serum depletion, we observed that the localization of CP110 and CEP97 to centrioles and centriolar satellites was dynamic: one hour after initiating ciliogenesis, CP110 and CEP97 at satellites decreased and, by 24 h, CP110 and CEP97 were absent from the mother centriole (**Figure 8A, B**).

CP110 interacts with satellite protein CEP290 (***Tsang et al., 2008***), so we hypothesized that CEP290 may hold CP110 at the satellites. Consistent with this model, CP110 no longer localized to satellites in cycling RPE1 cells upon CEP290 knockdown (**Figure 8- figure supplement 1C**). We propose that CP110 and CEP97 are centriolar satellite cargos which are wicked away from mother centrioles by centriolar satellites during early ciliogenesis.

Where does this overabundant CP110 and CEP97 accumulate? Using immunofluorescence microscopy of cycling cells treated with nocodazole, we examined the localization of CP110 and CEP97 to centrioles. In the absence of PCM1, CP110 and CEP97 over-accumulated at both centrioles (**Figure 8E-H**), suggesting that centriolar satellites restrict CP110 and CEP97 accumulation at centrioles.

We conclude that centriolar satellites restrict CP110 and CEP97 levels at centrioles, the removal of which promotes ciliogenesis in some cell types. Centriolar satellites help promote timely ciliary vesicle formation and remove CP110 and CEP97 from the mother centriole, enabling recruitment of IFT and construction of the transition zone, early steps in ciliogenesis important for the prevention of ciliopathy-associated phenotypes such as hydrocephaly (**Figure 9**).

**Figure 9.**
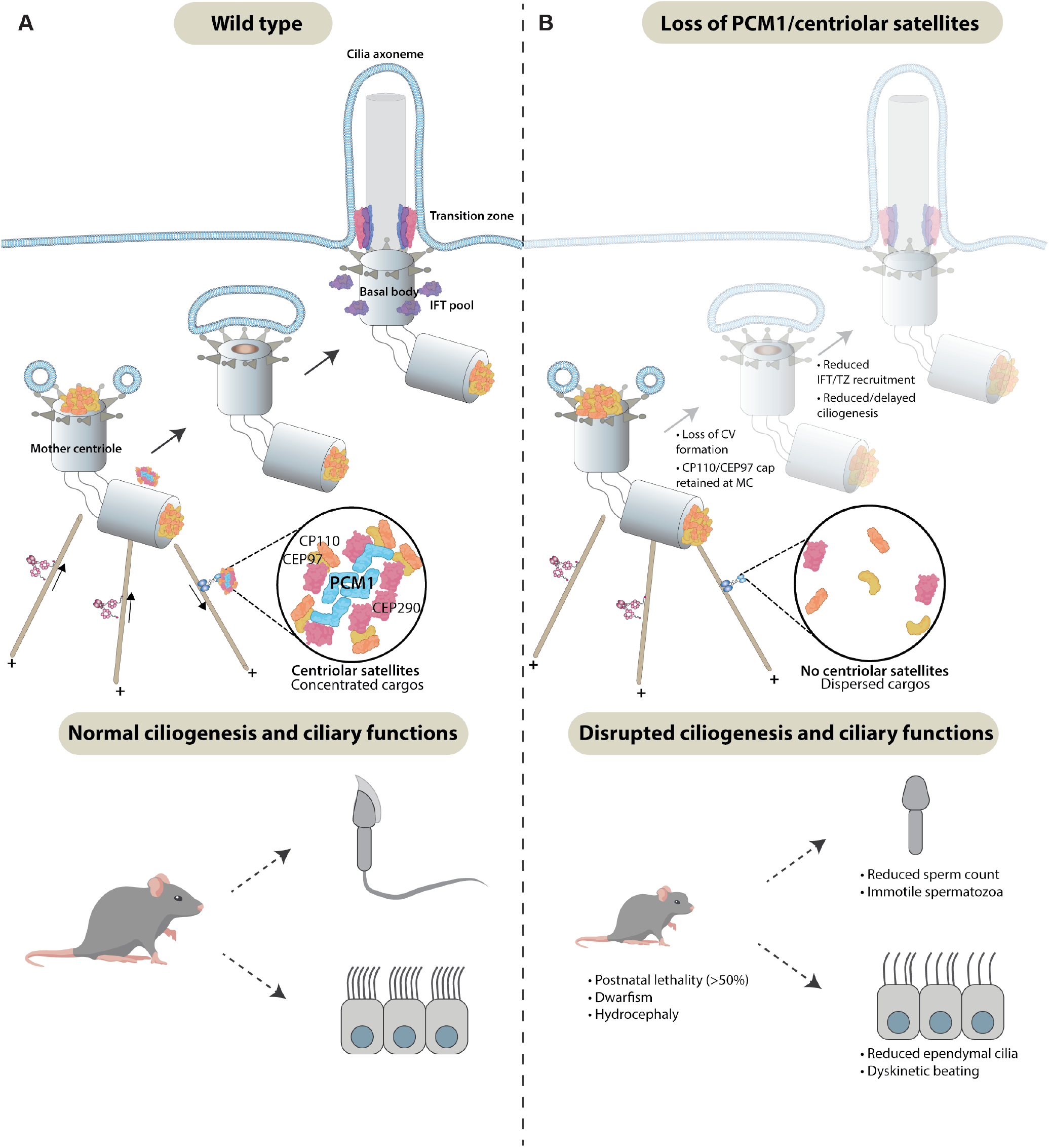
Centriolar satellites remodel centrioles to promote ciliogenesis. **(A)** PCM1 (cyan) scaffolds centriolar satellites, dynamic and heterogeneous condensates of centriolar proteins. During ciliogenesis, we propose that centriolar satellites remove, or wick away, CP110 and CEP97 from the mother centriole. Departure of CP110 and CEP97 is important for subsequent steps in ciliogenesis, including centriolar vesicle formation, transition zone formation and IFT-B recruitment. **(B)** In the absence of PCM1 and centriolar satellites, CP110 and CEP97 are not efficiently removed during ciliogenesis, disrupting subsequent steps, impeding ciliogenesis and leading to hydrocephaly and other ciliopathy-associated phenotypes.

## Discussion

### PCM1 is required for select cilia functions *in vivo*

Cilia are essential for key events in mammalian development; mice lacking cilia die during embryogenesis with developmental defects including randomized left-right axis specification and polydactyly (***Ferrante et al., 2006; Huangfu et al., 2003***). *Pcm1*^−/−^ mice survived at Mendelian ratios throughout gestation, displayed no evidence of complex cardiac defects associated with situs abnormalities, craniofacial anomalies or polydactyly. As PCM1 is not required for left-right axis, face or limb bud patterning in utero and is dispensable for ciliogenesis in primary MEFs, centriolar satellites are not required for mammalian ciliogenesis in many cell types.

Most *Pcm1*^−/−^ mice died perinatally with hydrocephaly, delayed formation and disrupted function of ependymal cilia, oligospermia and delayed tracheal epithelial cell ciliogenesis. In addition, *Pcm1*^−/−^ mice exhibited cerebellar hypoplasia and partially penetrant hydronephrosis, both of which can be caused by defective Hedgehog signaling, a signal transduction pathway dependent on cilia (***Huangfu et al., 2003; Spassky et al., 2008; Wallace, 1999; Wechsler-Reya and Scott, 1999; Yu et al., 2002***).

Recently, a mouse *Pcm1* gene trap was described (***Monroe et al., 2020***). Aged mice homozygous for this allele exhibited enlarged brain ventricles, progressive neuronal cilia maintenance defects and late-onset behavioral changes, but without perinatal lethality and other early cilia-associated phenotypes. While background differences may influence penetrance and expressivity, it is possible that the absence of reported hydrocephaly and other ciliopathy-related phenotypes indicates that the *Pcm1* gene trap allele is hypomorphic.

Most human ciliopathies affect select tissues (***Reiter and Leroux, 2017***). For many ciliopathies, it remains unclear why tissues are differentially sensitive to ciliary defects. As PCM1 is particularly required for mammalian cilia function in ependymal cells and sperm, differential requirements for centriolar satellite function may be one determinant of tissue specificity.

### PCM1 and centriolar satellites promote centriole amplification in ependymal cells

Centriole duplication is tightly controlled in cycling cells, restricting generation to two new centrioles per cell cycle (***Nigg and Holland, 2018***)N. In marked contrast, postmitotic multiciliated cells produce tens to hundreds of centrioles. This centriole amplification has been proposed to occur by two mechanisms; (i) generation of new centrioles in proximity to the parental centrioles and (ii) generation via deuterosomes, electron dense structures unique to multiciliated cells (***Mercey et al., 2019a; Nanjundappa et al., 2019; Zhao et al., 2013***,?). However, centriole amplification and multiciliogenesis are not blocked in the absence of deuterosomes or parental centrioles (***Mercey et al., 2019a***,b; ***Zhao et al., 2019***), indicating that a third mechanism may exist.

A previous study demonstrated that knockdown of *Pcm1* in cultured mouse ependymal cells did not affect centriole number, but did alter ciliary structure (***Zhao et al., 2021***). We found that, in the absence of PCM1, ependymal cells displayed retarded centriole amplification and multiciliogenesis, which resulted in hydrocephaly. Our data indicate that PCM1, unlike deuterosomes, is critical for timely centriole amplification in ependymal and tracheal cells. We propose that PCM1 is key to this previously postulated third mechanism of centriole amplification.

In addition to delayed centriole amplification, ependymal cells lacking PCM1 generated extremely elongated (3-7 *μ*m) centriole-related structures containing FOP and Centrin2. These centriolerelated structures were present within the cytoplasm, distant from the apical domain where basal bodies support ciliogenesis and support a role for PCM1 in controlling centriole morphology.

### Centriolar satellites promote the timely removal of CP110 and CEP97 to support ciliogenesis

Our work indicates that PCM1 and centriolar satellites help control the composition of centrioles. We found that, in MEFs, RPE1 and ependymal cells, PCM1 promotes the removal of CP110 from distal mother centrioles, an early step in ciliogenesis (***Spektor et al., 2007***). Similarly, PCM1 restricts levels of CEP131, CEP290 and CEP97 at centrioles. Recent work showed that *Pcm1* knockdown in ependymal cells similarly increased CEP135 and CEP120 localization to basal bodies (***Zhao et al., 2021***). Thus, centriolar satellites restrict the centriolar accumulation of multiple proteins.

A previous study proposed a role for PCM1 in protecting Talpid3 from degradation by sequestering the E3 ligase, MIB1 away from the centrioles (***Wang et al., 2016***). We found that, in the absence of PCM1, MIB1 no longer localizes to centrioles but Talpid3 levels on *PCM1*^−/−^ centrioles were comparable to control centrioles, suggesting that PCM1 is not a critical determinant of Talpid3 levels. Talpid3 is required for distal appendage assembly and removal of daughter centriole proteins (e.g., Centrobin) from mother centrioles (***Wang et al., 2018***). We found that PCM1 is dispensable for distal appendage assembly and removal of Centrobin from the mother centrioles, further suggesting that PCM1 and centriolar satellites are not required for Talpid3-dependent functions. Thus, centriolar satellites limit the centriolar localization of some, but not all, centriole components.

In the absence of PCM1, total cellular CP110 levels are elevated and CP110 and CEP97 levels are elevated at centrioles, indicating a role for centriolar satellites in CP110 degradation. As we observe CP110 and CEP97 transiently at satellites, and confirm by co-immunoprecipitation recent proteomic studies identifying CP110 and CEP97 as potential satellite constituents (***Gheiratmand et al., 2019; Quarantotti et al., 2019***), we propose that satellites transport CP110 and CEP97 away from centrioles for degradation, presumably by proteosomes. Previous identification of changes in centriolar satellite composition and distribution in response to environmental cues and stressors may indicate that satellites help dispose of proteins beyond CP110 and CEP97 (***Joachim et al., 2017; Prosser and Pelletier, 2020; Tollenaere et al., 2015; Villumsen et al., 2013***).

The transient localization of CP110 to centriolar satellites is dependent on its interactor, CEP290. As inhibition of ciliogenesis by CP110 is dependent on CEP290 (***Tsang et al., 2008***), we suggest that one function for CEP290 may be to recruit CP110 to satellites for removal from mother centrioles.

In vertebrates, CP110 is required for docking of the mother centriole to preciliary vesicles (***Walentek et al., 2016; Yadav et al., 2016***) and is removed from the mother centriole subsequent to docking, suggesting that it has both positive and inhibitory roles in ciliogenesis (***Lu et al., 2015***). Our finding that PCM1 promotes both CP110 removal and vesicular docking of the mother centriole suggests that both processes are intimately connected. However, PCM1 and centriolar satellites may contribute to preciliary vesicle docking through independent mechanisms. For example, although we demonstrate that PCM1 is dispensable for distal appendage formation, the subtle differences in some distal appendage component levels in *PCM1*^−/−^ cells could manifest as changes to distal appendage composition or conformation in ways that compromise vesicle docking.

Interestingly, centriolar satellites promote removal of CP110 from both RPE1 cells and MEFs, but are dispensable for ciliogenesis in MEFs, revealing that removal of all CP110 from mother centrioles is not a precondition for ciliogenesis in some cell types. Multiple roles for CP110, both promoting and inhibiting ciliogenesis, have previously been described (***Gonçalves et al., 2021; Walentek et al., 2016; Yadav et al., 2016***). One possible explanation for the cell-type specificity of PCM1 function is that centriolar satellites remove CP110 from mother centrioles in all cell types, but different thresholds of CP110 reduction are required to initiate ciliogenesis in different cell types. Perhaps CP110 removal from mother centrioles is especially important for cells, like many epithelial cells, in which basal bodies dock directly to the plasma membrane, rather than to a ciliary vesicle.

Thus, unlike core centriolar proteins, some of which are trafficked via centriolar satellites, centriolar satellites themselves are not essential for all centriole-and cilium-dependent events in many mammalian cell types as the threshold to which different cells in vivo tolerate disruption of cascades to remodel centrioles during cilia assembly or disassembly appears to be different. Perhaps in the crowded environment at the heart of the centrosome, diffusion is insufficient for the timely delivery and removal of centriolar proteins, and PCM1 and centriolar satellites promote centriole amplification and ciliogenesis by coupling assembly and/or degradation of centriolar components in the satellites to their active transport to and from centrioles on microtubules.

## Methods and Materials

### Generation of mouse models

Animals were maintained in SPF environment and studies carried out in accordance with the guidance issued by the Medical Research Council in “Responsibility in the Use of Animals in Medical Research” (July 1993) and licensed by the Home Office under the Animals (Scientific Procedures) Act 1986 under project license number P18921CDE in facilities at the University of Edinburgh (PEL 60/6025). *Pcm1* null mice (*Pcm1*^Δ5−14^*/* : *Pcm1*^*em*1*P mi*^ MGI:6865681 and *Pcm1*^Δ796−800^: *Pcm1*^*em*2*P mi*^ MGI:6865682) were generated using CRISPR/Cas9 as described in **Figure 1- figure supplement 1**, using guides detailed in **Table 1**. Genotyping was performed using primers detailed in **Table 2** followed by Sanger sequencing (for *Pcm1*^Δ5−14^ or digestion with DdeI (for *Pcm1*^Δ796−800^, or alternately genotyping was performed by Transnetyx. *Pcm1*^*SNAP*^ animals were generated with CRISPR Cas9 targeting first coding exon 2 (**Table 1**) and a SNAP tag was inserted after the ATG, followed by a GSGG linker, using a repair template with 700 nt homology arms, detailed in **Table 1**, resulting in a gene encoding N-terminally SNAP tagged PCM1 in the endogenous locus. Genotyping was performed using primers detailed in **Table 2** or alternately by Transnetyx.

**Table 1.**
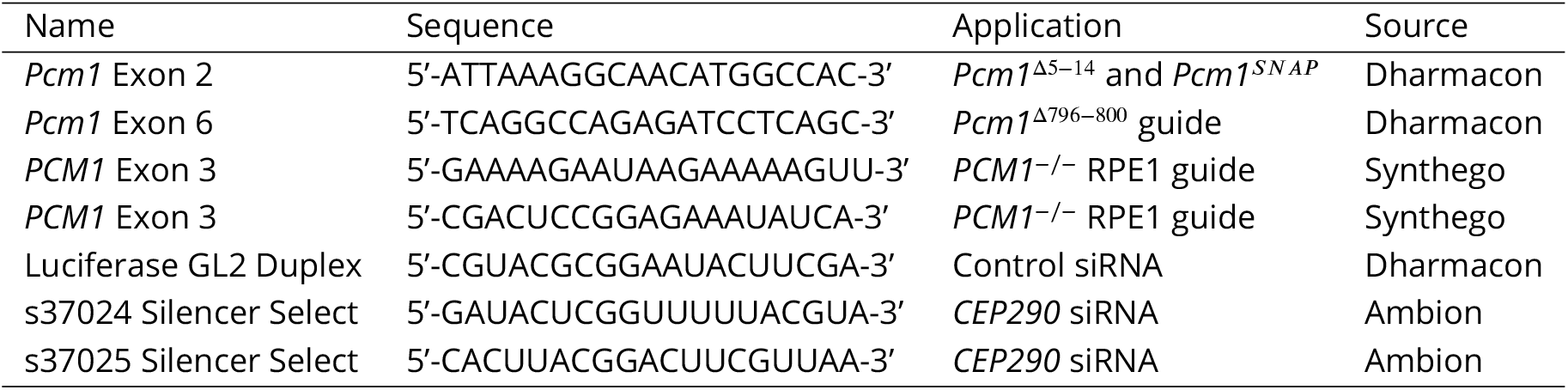
CRISPR guides and siRNAs.

**Table 2.**
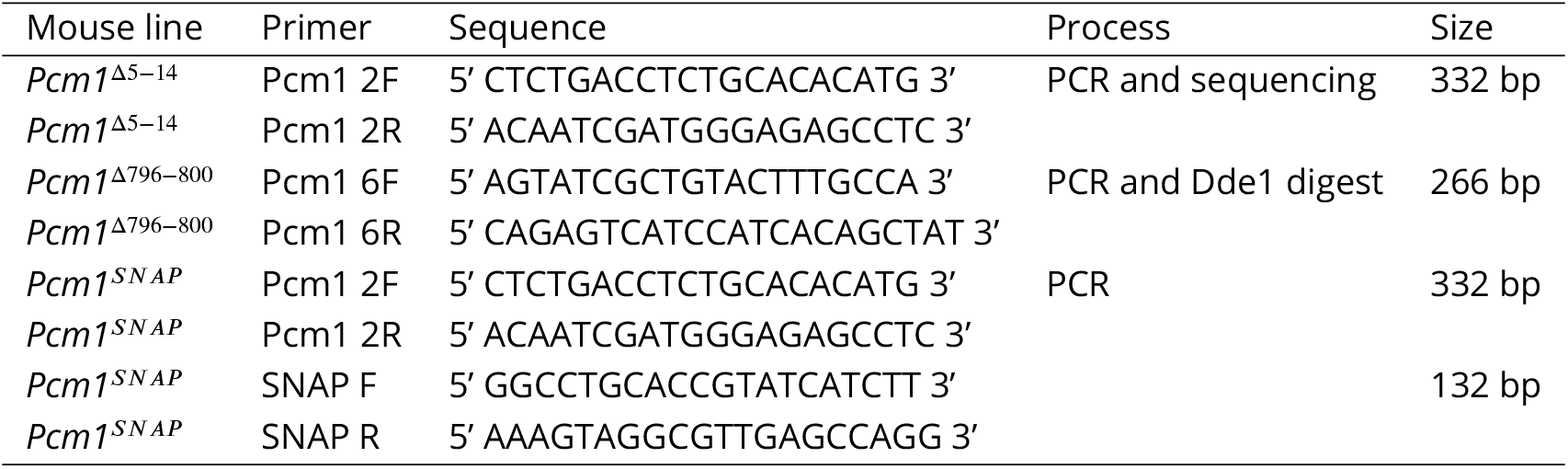
Genotyping primers.

### Mouse gait analysis

Gait analysis was performed on a CatwalkTM XT according to manufacturer’s instructions. Briefly, mice were habituated to the Catwalk for 5 min, and then the glass was cleaned prior to acquisition. Each mouse (n>4 per experimental group) was then allowed to perform at least 3 runs across the Catwalk, which records paw position and analyses gait patterns using the Catwalk XT 10.6 Acquisition and Analysis Software.

### Retinal imaging

Electroretinograms and fundal imaging was performed as described in (***Findlay et al., 2018***). PCM1-SNAP retinal labelling was carried out under inhaled anesthesia. 1.5 *μ*l of 0.6 *μ*M SNAP-Cell 647-SiR (New England Biolabs) was injected into the mouse vitreous under direct visualization using a Zeiss operating microscope. After 2 h, mice were sacrificed by cervical dislocation and eyes enucleated. Keratectomy, sclerectomy and lensectomy were performed and whole retinas isolated. Flat mount petaloid retinal explants were made and mounted, photoreceptor side up, on Menzel Glaser Superfrost Plus Gold slides (ThermoFisher Scientific; K5800AMNZ72). Nuclei were stained with DAPI and mounted in Prolong Gold under coverslip. Slices were imaged on an Andor Dragonfly spinning disc confocal.

### Cell lines and cell culture

Mouse embryonic fibroblasts (MEFs) were maintained as previously published (***Hall et al., 2013***). SNAP labelling was performed as previously described (***Quidwai et al., 2021***). Ependymal cells were isolated and cultured as published in (***Delgehyr et al., 2015***), and imaged 14-20 days post serum withdrawal. MTECs were isolated and cultured as described in (***Eenjes et al., 2018; You et al., 2002***). RPE1-hTERT (female, human epithelial cells immortalized with hTERT, Cat. No. CRL-4000) from ATCC were grown in DMEM (Life Technologies) or DMEM/F12 (ThermoFisher Scientific, 10565042) supplemented with 10% FBS at 37 °C with 5% CO_2_. *PCM1*^−/−^ RPE1 cells were generated as described previously (***Kumar et al., 2021***) (all figures except Figure 7 Supplement 1, in which case they were generated as in ***Gheiratmand et al***. (***2019***). Two *PCM1*^−/−^ RPE1 cell lines were generated using single guide RNAs (Table 1). Loss of PCM1 was confirmed by genotyping, immunoblotting, and immunofluorescence. Monoclonal *PCM1*^−/−^ RPE1 cell lines stably expressing eGFP or eYFP-PCM1 (plasmid a gift from Bryan Dynlacht; ***Wang et al***. (***2016***)) were generated using lentiviruses and manually selected based on fluorescence. To synchronize cells in G1/S aphidicolin (Sigma) was added to the culture medium at 2 *μ*g/ml for 16 h. To arrest cells in mitosis, taxol (paclitaxel; Millipore-Sigma) was added to the culture medium at 5 *μ*M for 16 h prior to rounded up cells being collected by mitotic shake-off. For arrest in G0, cells were washed 2x with PBS (Gibco) and 1x with DMEM (without serum) before being cultured in serum-free DMEM for 16 h. To disrupt cytoplasmic microtubules, cells were treated with 20 *μ*M nocodozole (Sigma, SML1665) for 1-2 h prior to fixation.

### RNA-mediated interference

hTERT RPE-1 cells were transfected with 20 nM (final concentration) of the respective siRNA for 48 h using Lipofectamine RNAiMAX (Invitrogen) according to the manufacturer’s instructions. Effective knockdown was confirmed by immunofluorescence microscopy. Details of individual siRNAs are provided in the **Table 1**.

### Total proteomics

MTECs were lysed in 0.1 % SDS in PBS plus 1X HALT protease inhibitor ThermoFisher Scientific, 78443), then processed by a multi-protease FASP protocol as described (***Wiśniewski and Mann, 2012***). In brief, SDS was removed and proteins were first digested with Lys-C (Wako) and subsequently with Trypsin (Promega) with an enzyme to protein ratio (1:50). 10 *μ*g of Lys-C and Trypsin digests were loaded separately and desalted on C18 Stage tip and eluates were analyzed by HPLC coupled to a Q-Exactive mass spectrometer as described previously (***Farrell et al., 2014***). Peptides and proteins were identified and quantified with the MaxQuant software package, and label-free quantification was performed by MaxLFQ (***Cox et al., 2014***). The search included variable modifications for oxidation of methionine, protein N-terminal acetylation, and carbamidomethylation as fixed modification. Peptides with at least seven amino acids were considered for identification. The false discovery rate, determined by searching a reverse database, was set at 0.01 for both peptides and proteins. All bioinformatic analyses were performed with the Perseus software ***Tyanova et al***. (***2016***)). Intensity values were log-normalized, 0-values were imputed by a normal distribution 1.8 *π* down of the mean and with a width of 0.2 *π*.

Proteomic expression data was analyzed in R (3.6.0) with the Bioconductor package DEP (1.6.1) (***Zhang et al., 2018***). To aid in the imputation of missing values only those proteins that are identified in all replicates of at least one condition were retained for analysis. The filtered proteomic data was normalized by variance stabilizing transformation. Following normalization, data missing at random, such as proteins quantified in some replicates but not in others, were imputed using the k-nearest neighbour approach. For differential expression analysis between the wildtype and mutant groups, protein-wise linear models combined with empirical Bayes statistics were run using the Bioconductor package limma (3.40.6) (***Ritchie et al., 2015***). Significantly differentially expressed proteins were defined by an FDR cutoff of 0.05. Total proteomic data are available via ProteomeXchange with identifier PXD031920 and are summarized in **Table 5**.

### Immunoblotting

Testes were lysed in RIPA buffer (Pierce) plus HALT protease inhibitor (ThermoFisher Scientific), homogenized with an electronic pestle for 1 minute, incubated at 4 °C with agitation for 30 min, sonicated for 3X 30 sec, and then clarified at 14,000 g at 4 °C for 20 min. RPE lysates were collected in 2x SDS-PAGE buffer and treated with benzonase nuclease (Millipore-Sigma) for 5 min. Samples were loaded into NuPAGE precast gels, transferred onto PVDF membrane (Amersham Hybond P, Cytiva), and then rinsed in water then TBST, and then blocked in 5% milk in TBS plus 0.1% Tween. Membranes were then incubated overnight at 4 °C in primary antibodies (**Table 3)** diluted in 5% milk TBST. 5 Membranes were then washed 3 × 10 min TBST, incubated in HRP conjugated secondary antibodies detailed in **Table 4** for 1 h at room temperature and developed using Pierce SuperSignal Pico Plus (Pierce) or ECL (GE Healthcare) reagent and imaged on ImageQuant.

**Table 3.**
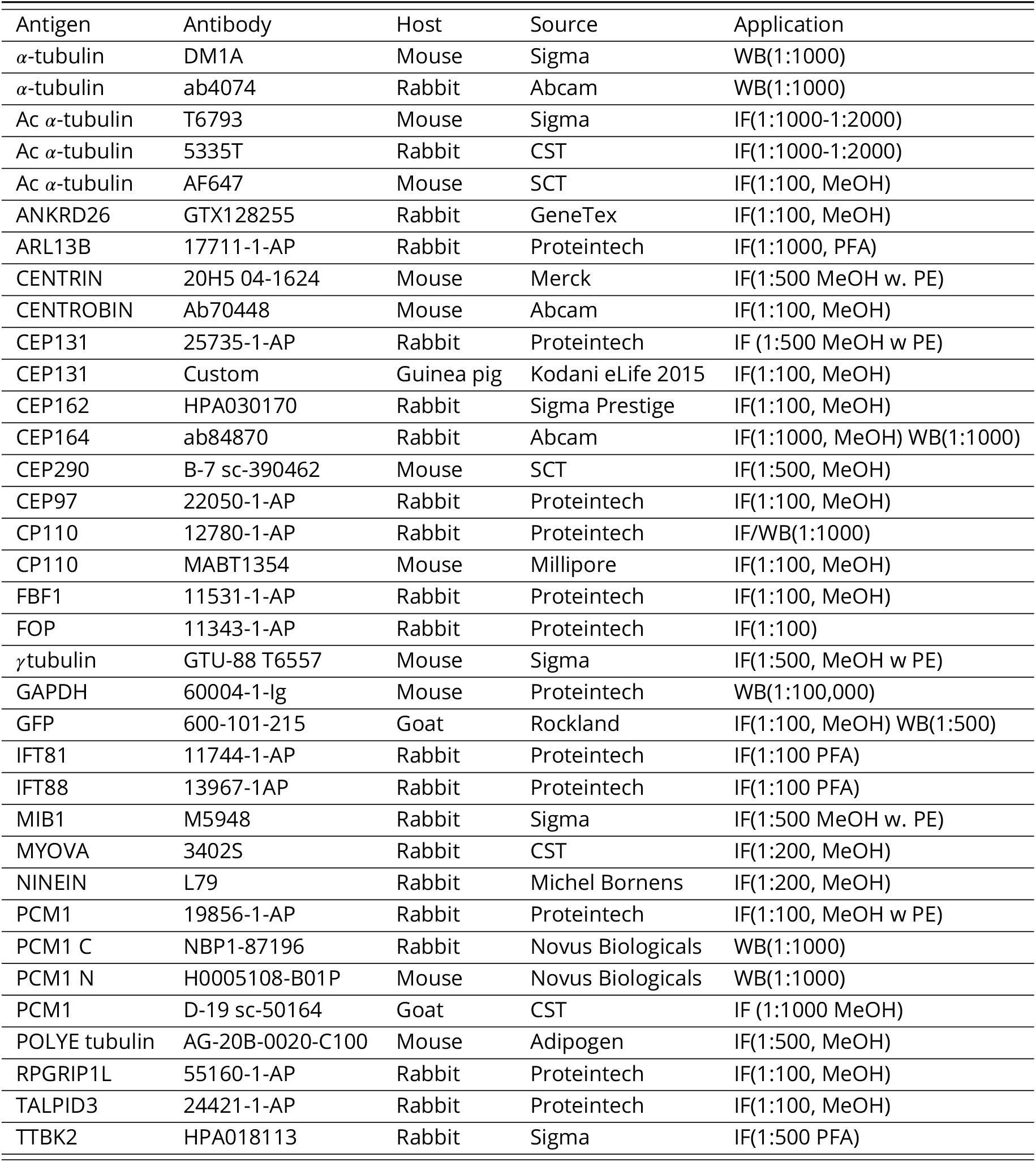
List of primary antibodies.

| Antigen | Antibody | Host | Source | Application |
| --- | --- | --- | --- | --- |
| α-tubulin | DM1A | Mouse | Sigma | WB(1:1000) |
| α-tubulin | ab4074 | Rabbit | Abcam | WB(1:1000) |
| Ac α-tubulin | T6793 | Mouse | Sigma | IF(1:1000-1:2000) |
| Ac α-tubulin | 5335T | Rabbit | CST | IF(1:1000-1:2000) |
| Ac α-tubulin | AF647 | Mouse | SCT | IF(1:100, MeOH) |
| ANKRD26 | GTX128255 | Rabbit | GeneTex | IF(1:100, MeOH) |
| ARL13B | 17711-1-AP | Rabbit | Proteintech | IF(1:1000, PFA) |
| CENTRIN | 20H5 04-1624 | Mouse | Merck | IF(1:500 MeOH w. PE) |
| CENTROBIN | Ab70448 | Mouse | Abcam | IF(1:100, MeOH) |
| CEP131 | 25735-1-AP | Rabbit | Proteintech | IF (1:500 MeOH w PE) |
| CEP131 | Custom | Guinea pig | Kodani eLife 2015 | IF(1:100, MeOH) |
| CEP162 | HPA030170 | Rabbit | Sigma Prestige | IF(1:100, MeOH) |
| CEP164 | ab84870 | Rabbit | Abcam | IF(1:1000, MeOH) WB(1:1000) |
| CEP290 | B-7 sc-390462 | Mouse | SCT | IF(1:500, MeOH) |
| CEP97 | 22050-1-AP | Rabbit | Proteintech | IF(1:100, MeOH) |
| CP110 | 12780-1-AP | Rabbit | Proteintech | IF/WB(1:1000) |
| CP110 | MABT1354 | Mouse | Millipore | IF(1:100, MeOH) |
| FBF1 | 11531-1-AP | Rabbit | Proteintech | IF(1:100, MeOH) |
| FOP | 11343-1-AP | Rabbit | Proteintech | IF(1:100) |
| γtubulin | GTU-88 T6557 | Mouse | Sigma | IF(1:500, MeOH w PE) |
| GAPDH | 60004-1-Ig | Mouse | Proteintech | WB(1:100,000) |
| GFP | 600-101-215 | Goat | Rockland | IF(1:100, MeOH) WB(1:500) |
| IFT81 | 11744-1-AP | Rabbit | Proteintech | IF(1:100 PFA) |
| IFT88 | 13967-1AP | Rabbit | Proteintech | IF(1:100 PFA) |
| MIB1 | M5948 | Rabbit | Sigma | IF(1:500 MeOH w. PE) |
| MYOVA | 3402S | Rabbit | CST | IF(1:200, MeOH) |
| NINEIN | L79 | Rabbit | Michel Bornens | IF(1:200, MeOH) |
| PCM1 | 19856-1-AP | Rabbit | Proteintech | IF(1:100, MeOH w PE) |
| PCM1 C | NBP1-87196 | Rabbit | Novus Biologicals | WB(1:1000) |
| PCM1 N | H0005108-B01P | Mouse | Novus Biologicals | WB(1:1000) |
| PCM1 | D-19 sc-50164 | Goat | CST | IF (1:1000 MeOH) |
| POLYE tubulin | AG-20B-0020-C100 | Mouse | Adipogen | IF(1:500, MeOH) |
| RPGRIP1L | 55160-1-AP | Rabbit | Proteintech | IF(1:100, MeOH) |
| TALPID3 | 24421-1-AP | Rabbit | Proteintech | IF(1:100, MeOH) |
| TTBK2 | HPA018113 | Rabbit | Sigma | IF(1:500 PFA) |

**Table 4.** Secondary antibodies.

| Antigen | Host | Dilution | Source | Application |
| --- | --- | --- | --- | --- |
| ECL α-Mouse IgG, HRP-conj | Sheep | 1:7,500 | GE Healthcare | WB |
| α-Mouse IgG, HRP-conj | Goat | 1:5,000 | Jackson ImmunoResearch | WB |
| α-Mouse IgG H+L, HRP-conj | Goat | 1:5,000 | BioRad | WB |
| ECL α-Rabbit IgG, HRP-conj | Sheep | 1:10,000 | GE Healthcare | WB |
| α-Rabbit IgG, HRP-conj | Goat | 1:5,000 | Jackson ImmunoResearch | WB |
| α-Rabbit IgG H+L, HRP-conj | Goat | 1:5,000 | BioRad | WB |
| Alexa 488-conj-α-Mouse | Donkey | 1:500 | Invitrogen Molecular Probes | IF |
| Alexa 568-conj-α-Mouse | Donkey | 1:500 | Invitrogen Molecular Probes | IF |
| Alexa 594-conj-α-Mouse | Donkey | 1:500 | Invitrogen Molecular Probes | IF |
| Alexa 647-conj-α-Mouse | Donkey | 1:500 | Invitrogen Molecular Probes | IF |
| Alexa 488-conj-α-Mouse | Donkey | 1:500 | Invitrogen Molecular Probes | IF |
| Alexa 568-conj-α-Rabbit | Donkey | 1:500 | Invitrogen Molecular Probes | IF |
| Alexa 594-conj-α-Rabbit | Donkey | 1:500 | Invitrogen Molecular Probes | IF |
| Alexa 647-conj-α-Rabbit | Donkey | 1:500 | Invitrogen Molecular Probes | IF |
| Alexa 647-conj-α-Goat | Donkey | 1:500 | Invitrogen Molecular Probes | IF |
| Alexa 647-conj-α-Guinea pig | Goat | 1:500 | Invitrogen Molecular Probes | IF |

### Co-immunoprecipitation

Co-immunoprecipitation assays and western blots were performed as described previously (***Kumar et al., 2021***) using GFP trap magnetic agarose beads (Chromotek, gtma-10).

### Ventricle wholemount

Ventricles were dissected according to (***Mirzadeh et al., 2010a***), pre-extracted with 0.1% Triton X in PBS for 1 minute, then fixed in 4% PFA or ice cold methanol for at least 24 h at 4 °C, followed by permeabilization in PBST (0.5% Triton X-100) for 20 min room temperature. Ventricles were blocked in 10% donkey serum in TBST (0.1% Triton-X) or 4% BSA in PBST (0.25% Triton X-100) for one h at room temperature, then placed ependymal layer down in primary antibodies (**Table 3**) in 4% BSA PBST (0.25% Tween-20) or 1% donkey serum in TBST (0.1% Triton-X) for at least 12 h. Ventricles were washed in PBS 3 × 10 min and secondaries (**Table 4**) in 4% BSA in PBST (0.25% Triton X-100) or 1% donkey serum in TBST (0.1% Triton-X) were added at 4 °C for at least 12 h. Ventricles were washed in PBS 3 × 10 min, and mounted on glass bottom dishes (Nest, 801002) in Vectashield (VectorLabs), immobilized with a cell strainer (Greiner Bio-One, 542040).

### Histology

Kidneys and brains were fixed in 4% PFA/PBS, testes were fixed in Bouin’s fixative, and eyes were fixed in Davidson’s fixative according to standard protocols. Tissues were serially dehydrated and embedded in paraffin. Microtome sections of 8 *μ*m thickness were examined histologically via haematoxylin and eosin (HE) or periodic acid-Schiff (PAS) staining.

For immunofluorescent analysis, paraffin sections were dewaxed and re-hydrated via ethanol series, followed by antigen retrieval by boiling the sections for 15 min in the microwave in citrate buffer. Sections were blocked in 10% donkey serum/0.1% Triton-X100 in PBS and primary antibodies were diluted in 1% donkey serum/PBS (**Table 3**). Slides were washed and incubated in Alexafluor conjugated secondary antibodies (**Table 4**), washed and mounted in ProLong Gold (ThermoFisher Scientific).

### Immunofluorescence

MEFs were processed for immunofluorescence as published (***Hall et al., 2013***). Briefly, cells were washed twice with warm PBS, then fixed in either 4% PFA in 1X PHEM/PBS 10 min at 37 °C or preextracted for 30 sec on ice in PEM (0.1 M PIPES pH 6.8, 2 mM EGTA, 1 mM MgSO_4_ prior to fixing in ice cold methanol on ice for 10 min according to **Table 3**, then washed twice with PBS. Cells were permeabilized and blocked with 10% donkey serum in 0.1% Triton-X 100/TBS for 60 min at room temperature, or overnight at 4 °C. Primary antibodies (**Table 3**) were added to samples and incubated for 4 °C overnight, in dilutant made of 1% donkey serum in 0.1% Triton X-100/TBS. Samples were washed in 0.1% Triton-X 100/TBS 4-6 times, 10 min each. Secondary antibodies diluted in 1% donkey serum and 0.1% Triton X-100/TBS were added for 60 min at room temperature, in some cases co-stained with AlexaFluor 647 Phalloidin (ThermoFisher Scientific), added with the secondaries **Table 4** at 1/500 for 1 h at room temperature. Samples were washed with 0.1% Triton-X 100/TBS 4-6 times 10 min, stained with DAPI (1:1000) in 0.1% Triton X-100/TBS for 5 min at room temperature, and mounted using ProLong Gold antifade (ThermoFisher Scientific), according to the manufacturer’s instructions.

RPE1 cells were fixed with 100% cold methanol for 3 min and incubated in blocking buffer (2.5% bovine serum albumin (BSA), 0.1% Triton X-100 in PBS) for 1 h at room temperature (except in **Figure 7 – figure supplement 1**, where they were fixed in ice cold methanol for 10 min and incubated in 2% BSA in PBS for 10 min at room temperature). Coverslips were then incubated in primary antibodies (**Table 3**) in blocking buffer overnight at 4 °C or room temperature for 50 min, washed three times with PBS and incubated with secondary antibodies (**Table 4**) in blocking buffer for 1 h at room temperature along with Hoechst 33352 or DAPI (0.1 *μ*g/ml). Coverslips were washed three times with PBS and mounted with Prolong Diamond (ThermoFisher Scientific P36961) or ProLong Gold Antifade (Molecular Probes). For TTBK2 staining, cells were fixed with 4% PFA/PBS for 10 min in general tubulin buffer (80 mM PIPES, pH 7, 1 mM MgCl_2_, and 1 mM EGTA), permeabilized with 0.1% TX-100 and stained as described above (***Loukil et al., 2017***).

### Sperm preparation

Cauda and caput epididymides were dissected into M2 media (ThermoFisher Scientific). For live imaging, sperm were imaged in M2 media or 1% methyl cellulose (Sigma), in capillary tubes (Vitrotubes Mountain Leaks) sealed with Cristaseal (Hawskley). Sperm counts were performed on sperm from the cauda epididymides, diluted in water using a haemocytometer, only counting intact sperm (with both head and tail).

### Transmission electron microscopy

Samples were dissected into PBS. Samples were fixed in 2% PFA/2.5% glutaraldehyde/0.1 M Sodium Cacodylate Buffer pH 7.4 (Electron Microscopy Sciences). Lateral ventricle walls were fixed for 18 h at 4 °C othen subdissected into anterior, mid and posterior sections. Tissue was rinsed in 0.1 M sodium cacodylate buffer, post-fixed in 1% OsO_4_ (Agar Scientific) for one h and dehydrated in sequential steps of acetone prior to impregnation in increasing concentrations of resin (TAAB Lab Equipment) in acetone followed by 100%, placed in moulds and polymerized at 60 °C for 24 h.

Ultrathin sections of 70 nm were subsequently cut using a diamond knife on a Leica EM UC7 ultramicrotome. Sections were stretched with chloroform to eliminate compression and mounted on Pioloform filmed copper grids prior to staining with 1% aqueous uranyl acetate and lead citrate (Leica). They were viewed on a Philips CM100 Compustage Transmission Electron Microscope with images collected using an AMT CCD camera (Deben). RPE1 cells processed for TEM analysis were cultured on Permanox slides (Nunc 177445), serum starved for 1 h and processed as described previously (***Kumar et al., 2021***).

### Imaging

Brightfield images in **Figure 2** and **Figure 2- figure supplement 1** were imaged on a Hamumatsu Nanozoomer XR with x20 and x40 objectives. Macroscope images in **Figure 1** and **Figure 2** were imaged on a Nikon AZ100 Macroscope. Fluorescent images in **Figure 2, Figure 3- figure supplement 1, Figure 3- figure supplement 2, Figure 3- figure supplement 3** and **Figure 7- figure supplement 1** were taken on a Nikon A1+ Confocal with Oil 60 or 100x objectives with 405, Argon 561 and 640 lasers and GaSP detectors. Fluorescent images in **Figure 1, Figure 4A, 4B, 4I** and **Figure 4- figure supplement 1** were taken with Andor Dragonfly and Mosaic Spinning Disc confocal. Images in **Figure 3** and **Figure 6C** and **D** were taken with Nikon SORA with 405 nm 120 mW, 488 nm 200 mW and 561 nm 150 mW lasers, 100x 1.35 NA Si Apochromat objective and a Photometrics Prime 95B 11mm pixel camera. High speed video microscopy was performed on a Nikon Ti microscope with a 100X SR HP Apo TIRF Objective, and Prime BSI, A19B204007 camera, imaged at 250 fps. 3D-SIM imaging in **Figure 4E, F, H, Figure 5, Figure 5- figure supplement 1, Figure 6A, B, Figure 7** and **Figure 8** was performed using the GE Healthcare DeltaVision OMX-SR microscope equipped with the 60x/1.42 NA oil-immersion objective and three cMOS cameras. Immersion oil with refractive index of 1.518 was used for most experiments, and z stacks of 5-6 *μ*m were collected every 0.125 *μ*m. Images were reconstructed using GE Healthcare SoftWorx 6.5.2 using default parameters. Images for quantifications were collected at the widefield setting using the same microscope.

**Figure 8 Supplement 1** was imaged using a DeltaVision Elite high-resolution imaging system equipped with a sCMOS 2048 × 2048 pixel camera (GE Healthcare). Z-stacks (0.2 *μ*m step) were collected using a 60x 1.42 NA plan apochromat oil-immersion objective (Olympus) and deconvolved using softWoRx (v6.0, GE Healthcare).

### Image analysis

Image analysis was performed in NIS Elements, FIJI (***Schindelin et al., 2012***), QuPath (***Bankhead et al., 2017***), CellProfiler (***Stirling et al., 2021***) or Imaris. Macros used in this paper can be found on GitHub (https://github.com/IGC-Advanced-Imaging-Resource/Hall2022_Paper). Cerebellum and ventricle area was measured from PAS stained sagittal brain sections in QuPath. The number of ependymal cells with multiple basal bodies was calculated by segmenting FOP staining and cells in 2D using a CellProfiler pipeline. Briefly, an IdentifyPrimaryObjects module was used to detect the nuclei, followed by an IdentifySecondaryObjects module using the tubulin stain to detect the cell boundaries. Another Identify Primary objects module was used to detect the basal bodies and a RelateObjects module was used to assign parent-child relationships between the cells and basal bodies. The percentage of ciliated ependymal cells, and the number of ependymal cells with rosette-like FOP staining, and elongated FOP positive structures were counted by eye using NIS Elements Counts Tool. Analysis of cultured ependymal cells (beat frequency, number of cilia, coordinated beat pattern) was performed in FIJI by eye while blinded to genotype. CEP131 and MIB1 intensity at satellites was calculated in FIJI using a macro which segmented basal bodies with Gamma Tubulin, then drew concentric rings, each 0.5 *μ*m wider than the previous and calculated the intensity of MIB1 and CEP131 within these rings. CP110 intensity in MEFs was calculated by manually defining mother and daughter centrioles in FIJI, CP110 intensity in ependymals was calculated by segmenting FOP in 3D in Imaris and calculating CP110 intensity within this volume. Image quantification in RPE1 cells were performed using CellProfiler as described previously (***Kumar et al., 2021***). Images were prepared for publication using FIJI, Imaris, Adobe Photoshop, Illustrator and InDesign.

### Data analysis

Data analysis was carried out in Microsoft Excel, Graphpad Prism 6/9 and Matlab. Statistical tests are described in the Figure legends.

## Supporting information

Table 5. Differentially expressed proteins between wild type and Pcm1-/- mouse tracheal epithelial cells (mTECs) on at least one timepoint.

## Video Legends

- **Figure 2 – video 1. Wild-type sperm morphology and movement**.vSperm were isolated from wild-type testes and imaged in dilute methyl cellulose.
- **Figure 2 – video 2. *Pcm1***^−/−^ **sperm is immotile and lacks normal head structures**. Sperm were isolated from *Pcm1*^−/−^ testes and imaged in dilute methyl cellulose.
- **Figure 2 – video 3. *Pcm1***^−/−^ **sperm exhibit disrupted movement**. Sperm were isolated from *Pcm1*^−/−^ testes and imaged in media without methyl cellulose.
- **Figure 3 -video 1. Wild-type cultured ependymal cilia beat in a coordinated way**.
- **Figure 3 -video 2. *Pcm1***^−/−^ **ependymal cilia show uncoordinated, slow ciliary beat**.
- **Figure 3 -video 3.*Pcm1***^−/−^ **ependymal cilia show uncoordinated, slow ciliary beat**.
- **Figure 4 – video 1. Centriolar satellites frequently fuse and divide near the basal body**. PCM1-SNAP labelled with TMR-SNAP (yellow) in *Pcm1*^*SNAP*^ MEFs reveals that centriolar satellites show saltatory movement, coalescing and fragmenting around the base of cilia (SiR-tubulin, magenta). Scale bars represent 3 *μ*m.

## Acknowledgments

We thank the IGC Advanced Imaging Resource and the IGC Mass Spectrometry facility, as well as the Newcastle University Electron Microscopy Research services, and, in particular, Tracey Davey. Work by the group of JFR is supported by NIH R01GM095941, R01AR054396 and R01HD089918. DK was funded by the Jane Coffin Childs memorial foundation and Program for Biomedical Research award by the Sandler foundation. Work by the group of PM is supported by MRC intramural funding (MC UU 12018/26) and by the European Commission (H2020 Grant No. 866355). Work by the group of LP is supported by CIHR Foundation (FDN167279) and the Krembil Foundation. L.P is a Tier 1 Canada Research Chair in Centrosome Biogenesis and Function and S.L.P. was funded by a European Union Horizon 2020 Marie Skłodowska-Curie Global Fellowship (No. 702601).

## Author Contributions

EAH and DK developed the project, performed the bulk of the experiments, quantified and analyzed the results, prepared the figures and wrote the manuscript. EAH performed the mouse work and DK the work in RPE1 cells. SP performed some of the work on CP110 in RPE1 cells. PY performed some of the ependymal and tracheal wholemount experiments and helped with experimental design and manuscript proofreading. LR and LMK maintained mouse lines and helped with tissue processing and phenotyping. DD and PT helped with mTEC culture. VHP and JMGV performed TEM in Figure 5 and Figure 5- figure supplement 1. RM performed retinal SNAP labelling. LCM wrote image analysis scripts including analyzing satellite cargo intensity, quantifying ependymal basal body formation and CP110 intensity, quantifying cilia and centrosome number. MF performed analysis of brain histology. GG processed the mTEC total proteomic data. LW performed the MIB1 staining. TQ optimized SNAP staining in MEFs. LP advised on experimental design and proofread the manuscript. PM and JFR conceived of and contributed to the design of the project, advised on data analysis and presentation, and assisted in the writing of the manuscript.

## Declaration of Interests

The authors have declared no competing interests.

---

**Figure 1–Figure supplement 1.**
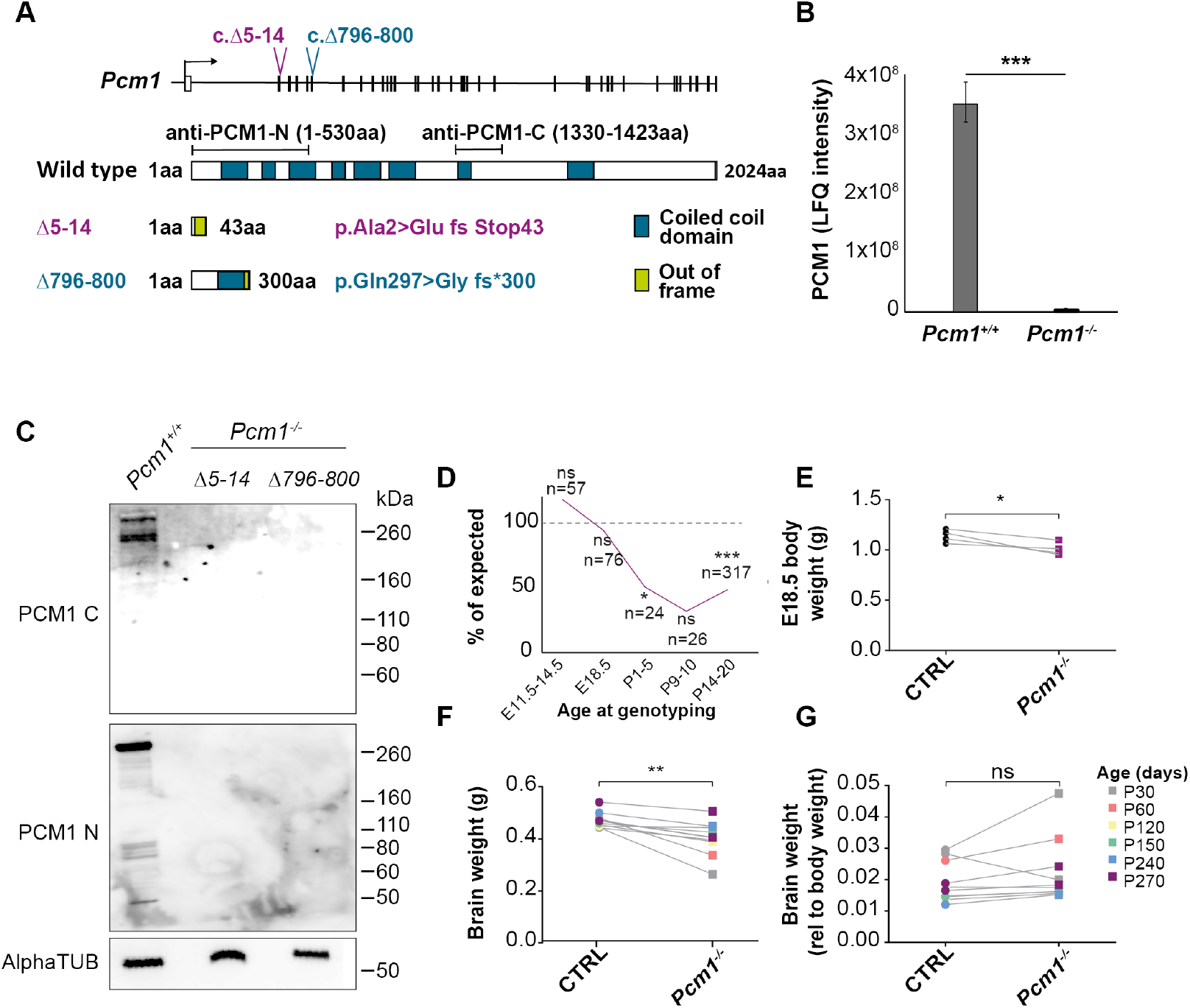
PCM1 promotes survival and growth. **(A)** Schematic of two *Pcm1* deletion mutations, *Pcm1*^Δ5−14^ and *Pcm1*^Δ796−800^, generated using CRISPR/Cas9. Both *Pcm1*^Δ5−14^ and *Pcm1*^Δ796−800^ create frameshifts. Schematic of PCM1 protein, indicating predicted coiled-coil domains and epitopes used for generating anti-PCM1 antibodies. **(B)** Label-free quantification (LFQ) mass spectrometric quantification of PCM1 in wild-type and *Pcm1*^−/−^ mouse tracheal epithelium cells (mTECs) differentiated for seven days in air-liquid interface. Student’s t-test ***P <0.01. **(C)** Immunoblot of *Pcm1*^Δ5−14^ and *Pcm1*^Δ796−800^ homozygous testis lysates for PCM1 using antibody directed against the PCM1 N-terminus (N), C-terminus (C) and *α*tubulin (loading control). **(D)** *Pcm1*^−/−^mice display partially penetrant perinatal lethality, with 50% of *Pcm1*^−/−^ pups lost just after birth. Graph shows number of *Pcm1*^−/−^ animals genotyped at each age as a percentage of the expected number, * P<0.05, *** P<0.001, Chi squared. n represents number of animals genotyped at given age. **(E)** E18.5 *Pcm1*^−/−^ embryos are smaller than their littermates. Each pair of points represents the average control or *Pcm1*^−/−^ null weight in a given litter, Paired t-test. *P<0.05, ***P<0.001. **F)** *Pcm1*^−/−^ brains are smaller than those of littermates. **(G)** The reduction of *Pcm1*^−/−^ brain size is proportional to the decrease in body weight. **(F, G)** Paired t-test, sex and litter matched. CTRL: heterozygote and wild-type measurements combined.

**Figure 1–Figure supplement 2.**
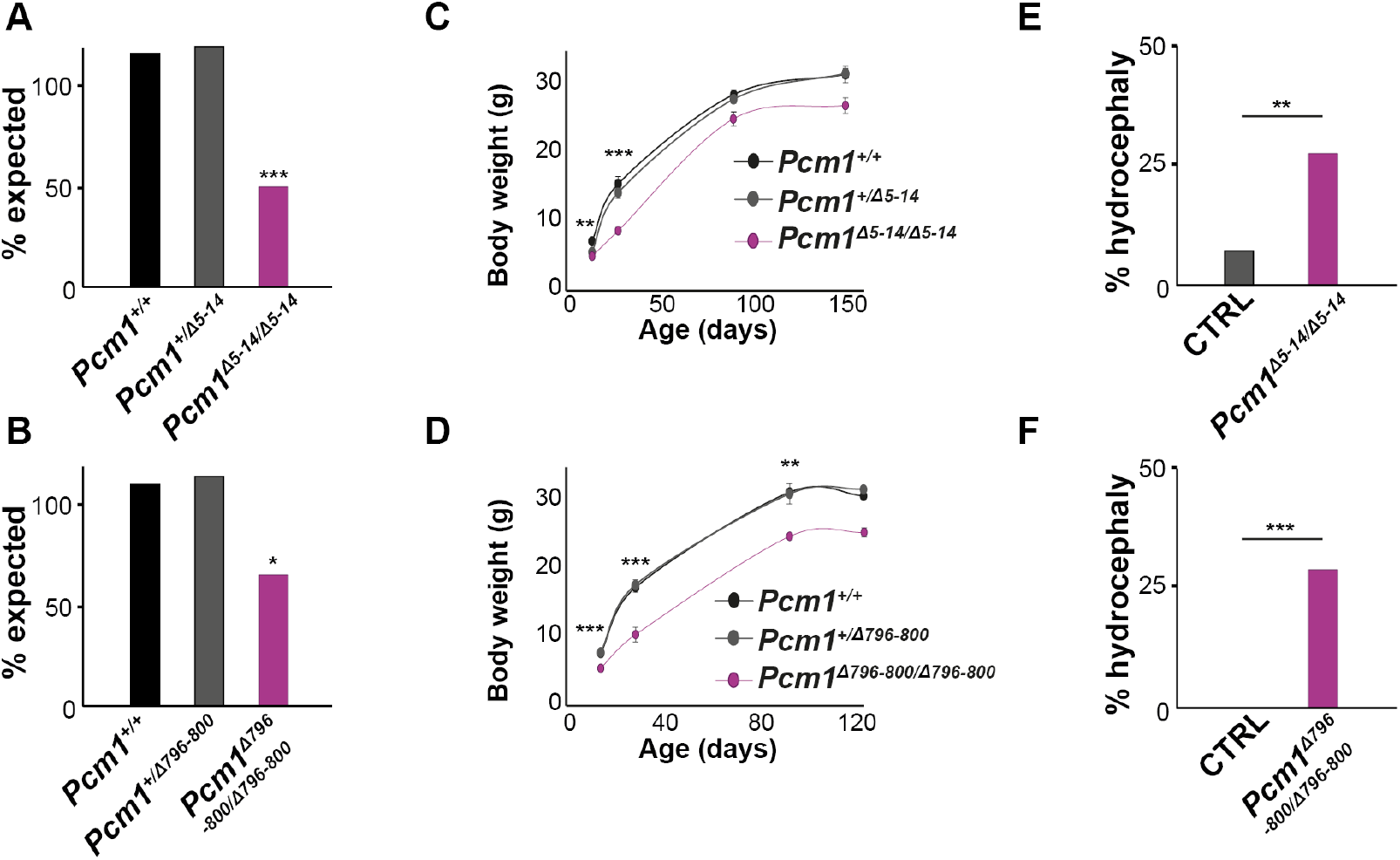
*Pcm1*^Δ5−14^ and *Pcm1*^Δ796−800^ mice exhibit comparable phenotypes. **(A, B)** *Pcm1*^Δ5−14^ homozygous and *Pcm1*^Δ796−800^ homozygous mice show comparable reductions in survival to genotyping. Chi squared: *** P< 0.01, *P< 0.05. **(C, D)** *Pcm1*^Δ5−14^ and *Pcm1*^Δ796−800^ homozygous mice show comparable reductions in body weight. Student’s t-test: ** P< 0.01, *** P< 0.001. **(E, F)** *Pcm1*^Δ5−14^ and *Pcm1*^Δ796−800^ homozygous mice show comparable incidences of overt hydrocephaly. Chi squared: ** P< 0.01, *** P< 0.001. CTRL: heterozygote and wild-type measurements combined.

**Figure 2–Figure supplement 1.**
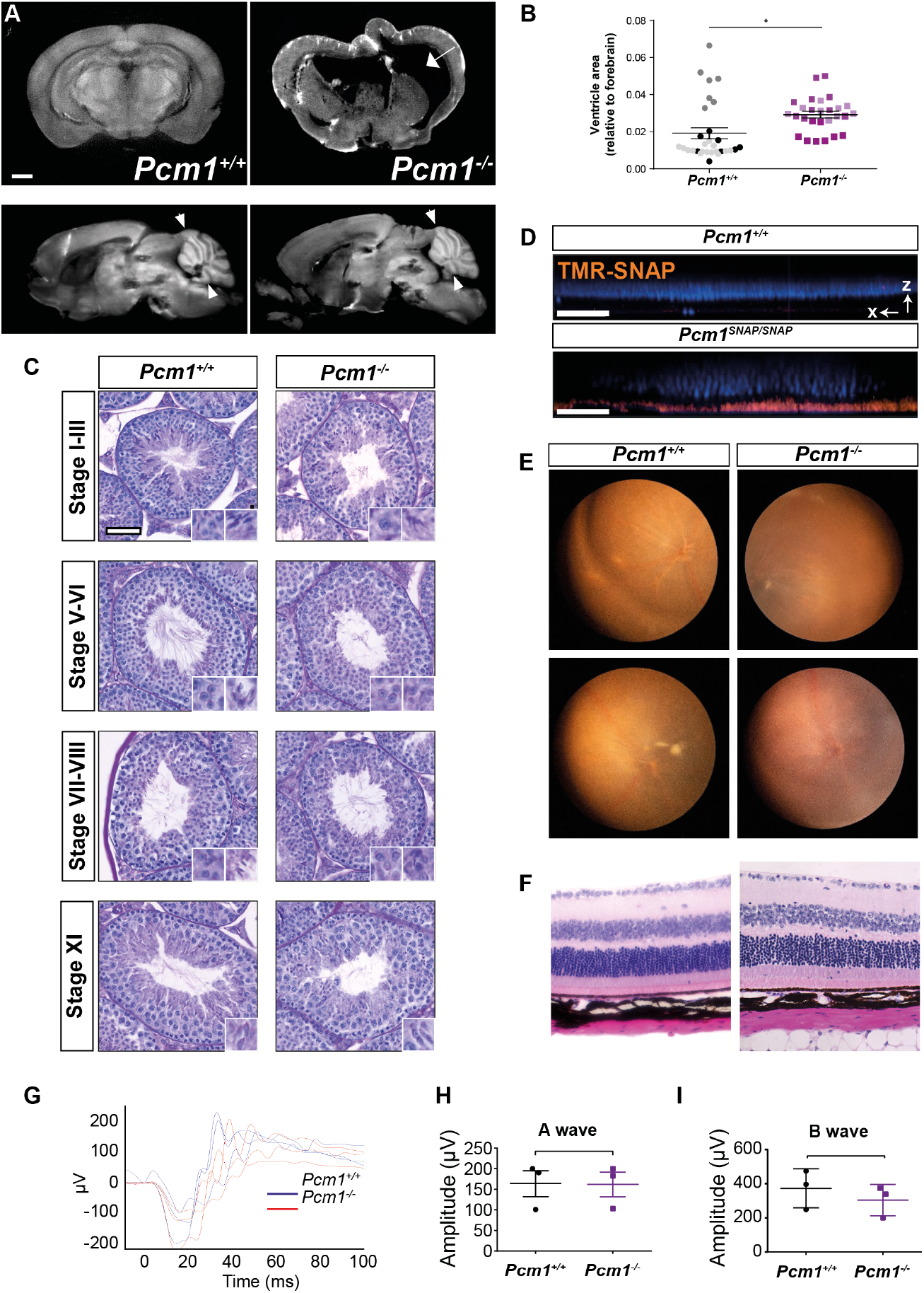
*Pcm1*^−/−^ mice display a subset of ciliopathy-associated phenotypes. **(A)** Coronal and sagittal optical sections from optical projection tomography (OPT) imaging of brains of *Pcm1*^+/+^ and *Pcm1*^−/−^ mice. The coronal image of the 6 week-old *Pcm1*^−/−^ brain depicts dilated ventricles (arrow). The sagittal section of the 8-month-old *Pcm1*^−/−^ brain depicts less severe hydrocephaly and cerebellar hypoplasia (arrowheads). Scale bar represents 1 mm. **(B)** Ventricle size relative to forebrain size of *Pcm1*^+/+^ and *Pcm1*^−/−^ mice without overt hydrocephaly exhibit dilatated ventricles. Each point depicts a measurement from an individual sagittal wax section, each shade represents a separate animal, *P<0.05, Student’s t-test. **(C)** PAS staining of stage-matched testis tubules. Insets show enlargements of developing spermatids. *Pcm1*^−/−^ tubules contain few sperm flagella. *Pcm1* null spermatids show abnormal development, from around Stage X, with abnormal manchette formation leading to hammerhead shaped sperm heads. Scale bar represents 50 *μ*m. **(D)** Intravitreous injection of TMR-SNAP into wild-type and *Pcm1*^−*SNAP* /*SNAP*^ eyes labels PCM1-SNAP in the retinas specifically of PCM1-SNAP-expressing animals. DAPI marks nuclei (blue). Displayed is a single X-Z slice from a confocal z stack. Scale bar represents 40 *μ*m. **(E)** Fundal imaging of 1-year-old wild type and *Pcm1*^−/−^ eyes. *Pcm1*^−/−^ mice do not display retinal degeneration by one year of age. **(F)** HE-stained sections of 13-month-old wild-type and *Pcm1*^−/−^ retinas. *Pcm1*^−/−^ eyes do not display histological signs of retinal degeneration. Scale bar represents 20 *μ*m. **(G)** Electroretinogram (ERG) of 9-month-old wild type and *Pcm1*^−/−^ mice. **(H)** Quantification of ERG A-waves, reflecting the rapid cornea negative potential. Student’s t-test. **(I)**. Quantification of ERG B-waves, reflecting the slow cornea positive potential. Student’s t-test.

**Figure 3–Figure supplement 1.**
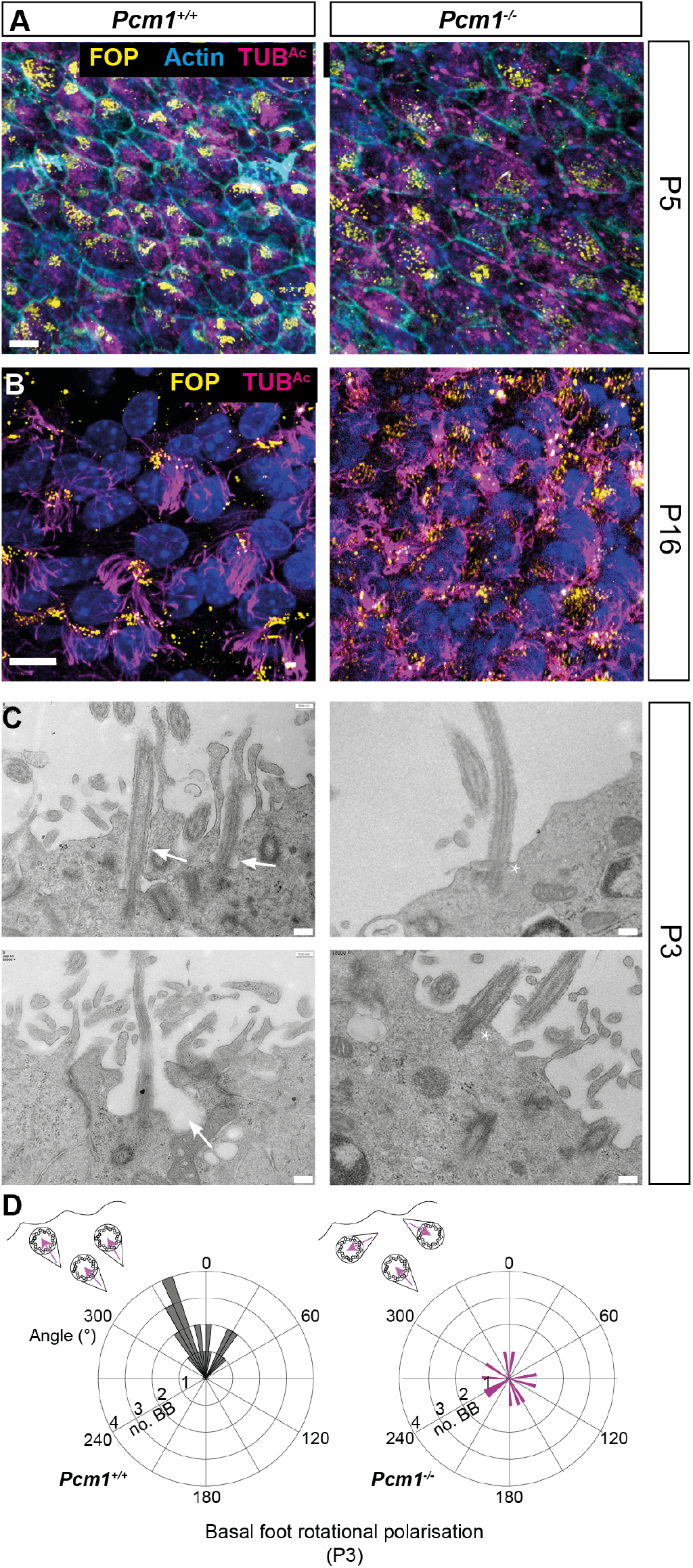
PCM1 is required for ciliary pocket formation and basal body polarization. **(A)** Ependymal cells from P5 wild-type and *Pcm1*^−/−^ ventricles immunostained for basal bodies (FOP, yellow), actin (phalloidin, cyan) and cilia (TUB^*Ac*^, magenta). At P5, this region of the *Pcm1*^−/−^ ventricles has equivalent numbers of cells with multiple basal bodies, but fewer of these ependymal cells have formed cilia. **(B)** P16 wild-type and *Pcm1*^−/−^ ependymal cells immunostained for basal bodies (FOP, yellow), cilia (TUB^*Ac*^, magenta), and nuclei (DAPI, blue). By P16, this region of the *Pcm1*^−/−^ ventricle has equivalent numbers of multiciliated cells. **(C)** TEM of P3 wildtype and *Pcm1*^−/−^ ependymal cilia. Wild-type cilia have ciliary pockets (arrows). *Pcm1*^−/−^ cilia do not (asterisks). **(D)** Rose plot of basal body rotational polarity quantified from TEM of P3 wild-type and *Pcm1*^−/−^ ependymal cells, angles plotted relative to the average angle of that micrograph, set at 0. Basal bodies are not planar polarized in *Pcm1*^−/−^ ependymal cells. Scale bars: 10 *μ*m (A and B) and 500 nm (C).

**Figure 3–Figure supplement 2.**
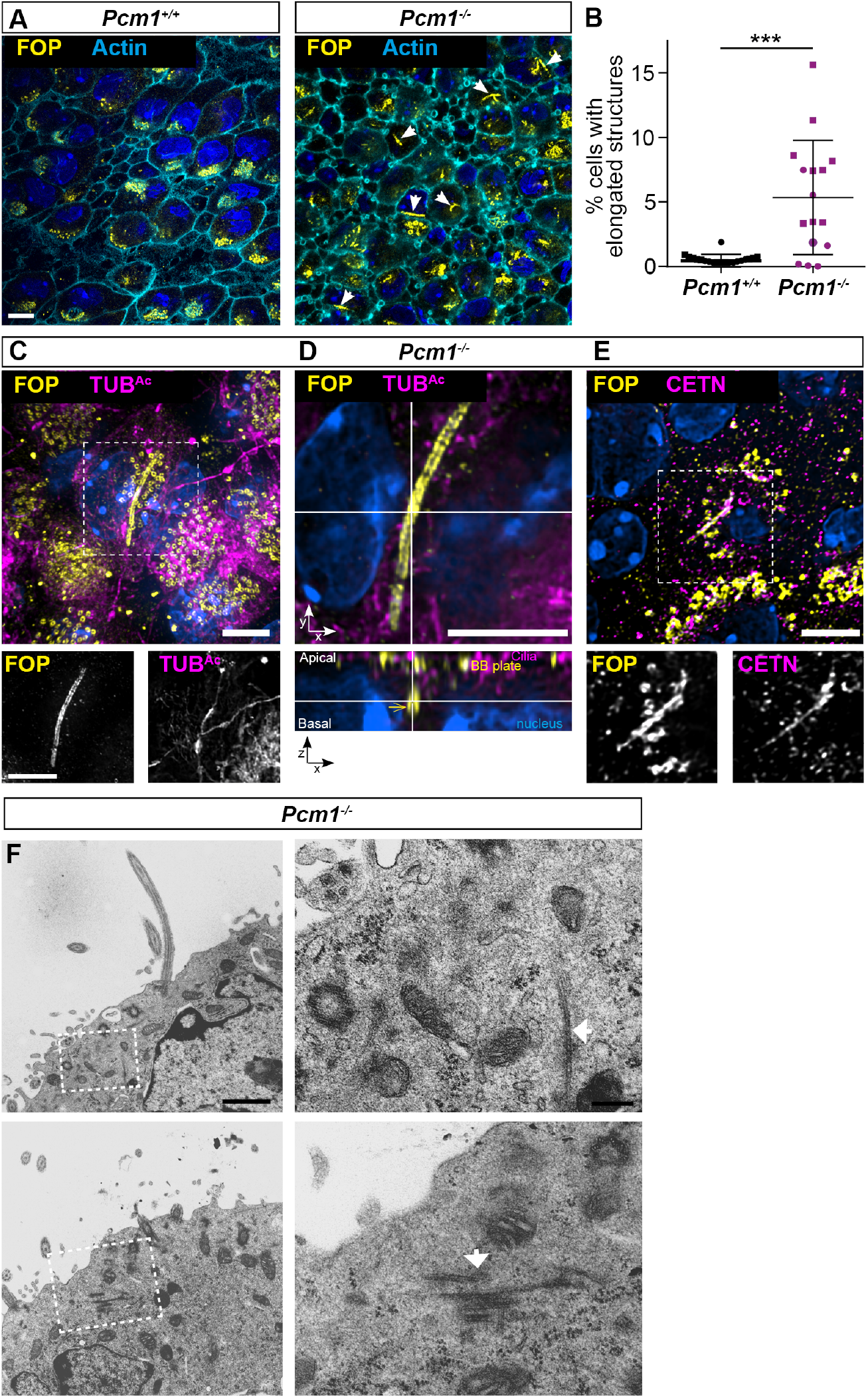
*Pcm1*^−/−^ ependymal cells form elongated centriole-like structures. **(A)** Ependymal cells from P3 wild-type and *Pcm1*^−/−^ ventricles immunostained for basal bodies (FOP, yellow), actin (phalloidin, cyan) and nuclei (DAPI, blue). A single optical section basal to the majority of basal bodies. *Pcm1*^−/−^ ependymal cells contain FOP-positive centriole-like structures (arrows) with a mean length of 5.0 *μ*m (standard deviation± 1.9 *μ*m). **(B)** Quantification of percentage of wild-type and *Pcm1*^−/−^ ependymal cells with elongated FOP-positive centriole-like structures. Student’s t-test, n = 2 *Pcm1*^+/+^ and 3 *Pcm1*^−/−^ mice, *** P<0.001. **(C)** P3 *Pcm1*^−/−^ ependymal cell immunostained for basal bodies (FOP, yellow), TUB^*Ac*^ (magenta) and nuclei (DAPI, blue), highlighting elongated centriole-like structures. Overlay with single channel images below. **(D)** Apical and lateral view of P3 *Pcm1*^−/−^ ependymal cell immunostained for basal bodies (FOP, yellow), TUB^*Ac*^ (magenta) and nuclei (DAPI, blue). **(E)** P3 *Pcm1*^−/−^ ependymal cell immunostained for basal bodies (FOP, yellow), Centrin (CETN, magenta) and nuclei (DAPI, blue). **(F)** TEM of ependymal cells from *Pcm1*^−/−^ P3 ventricle, highlighting elongated fibrillar structures (arrows) specific to mutant cells. Outlined areas on left are magnified on right. Scale bars: 10 *μ*m (A), 5 *μ*m (C-E), 1 *μ*m (F, left) and 200 nm (F, right).

**Figure 3–Figure supplement 3.**
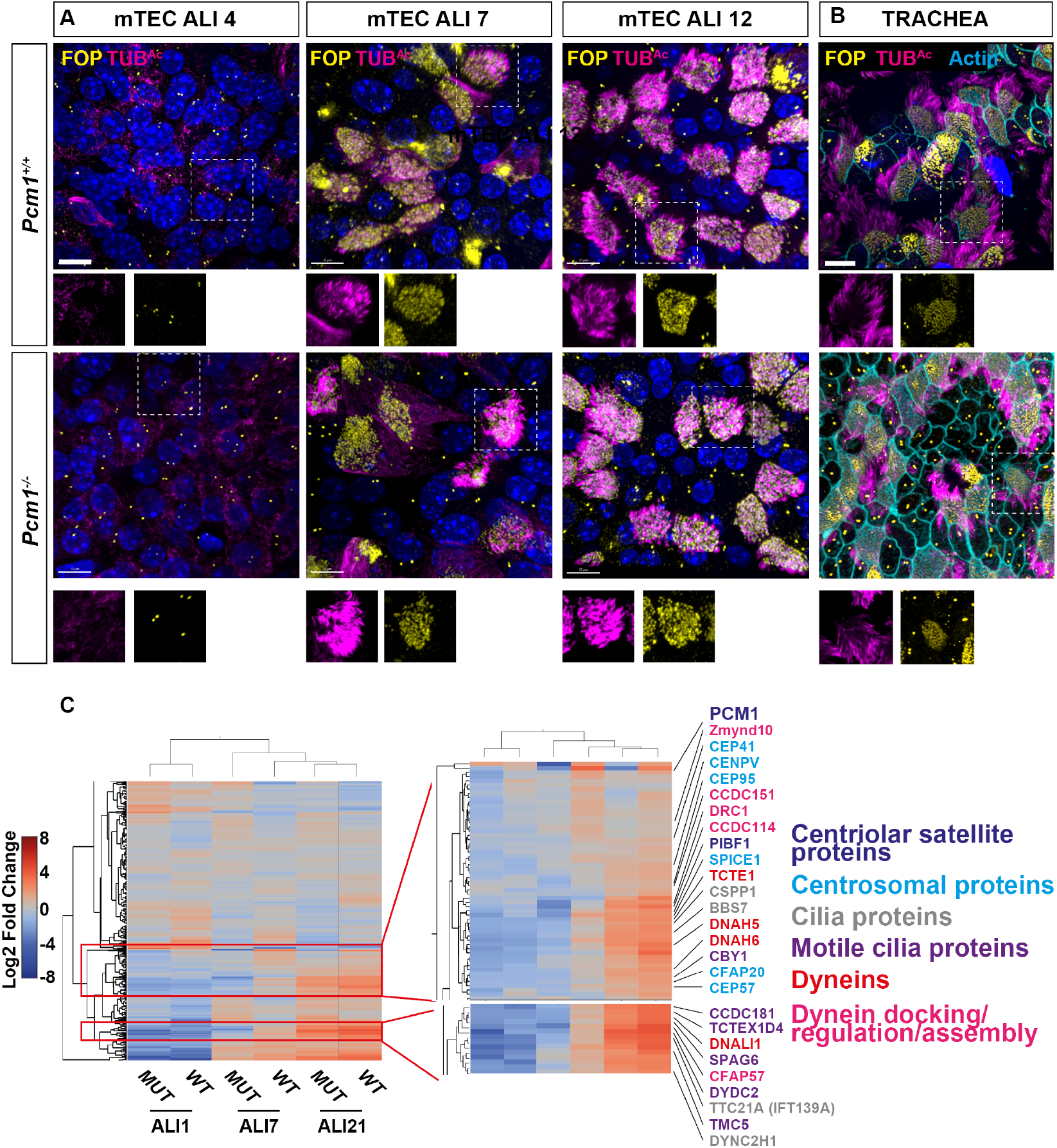
Delayed expression of ciliary proteins in *Pcm1*^−/−^ mTECs. **(A)** Wild-type and *Pcm1*^−/−^ mTECs immunostained for basal bodies (FOP, yellow), cilia (TUB^*Ac*^, magenta), and nuclei (DAPI, blue) 4, 7 or 12 days after placement at air-liquid interface (ALI). Below: magnifications of single channels from boxed areas. **(B)** Wild-type and *Pcm1*^−/−^ mTECs immunostained for basal bodies (FOP, yellow), cilia (TUB^*Ac*^, magenta), and actin (phalloidin, cyan). Scale bar represents 10 *μ*m. **(C)** Heatmap of total proteomic label-free quantification (LFQ) mass spectrometry analysis of wild-type and *Pcm1*^−/−^ mTECs, depicting all changed proteins (between timepoints and/or genotypes) at ALI D0 (unciliated), D7 (ciliating) and D21 (ciliated). Right: expansion of two clusters of proteins reduced in *Pcm1*^−/−^ mTECs at ALI D7. Font color indicates ontology: centriolar satellite proteins are in dark blue, centrosomal proteins are in light blue, ciliary proteins are in grey and ciliary motility proteins are in purple, including dyneins (red) and dynein docking/assembly factors (pink). The delayed induction of cilia-associated proteins in *Pcm1*^−/−^ mTECs suggests that, in the absence of PCM1, the ciliogenic program is retarded.

**Figure 4–Figure supplement 1.**
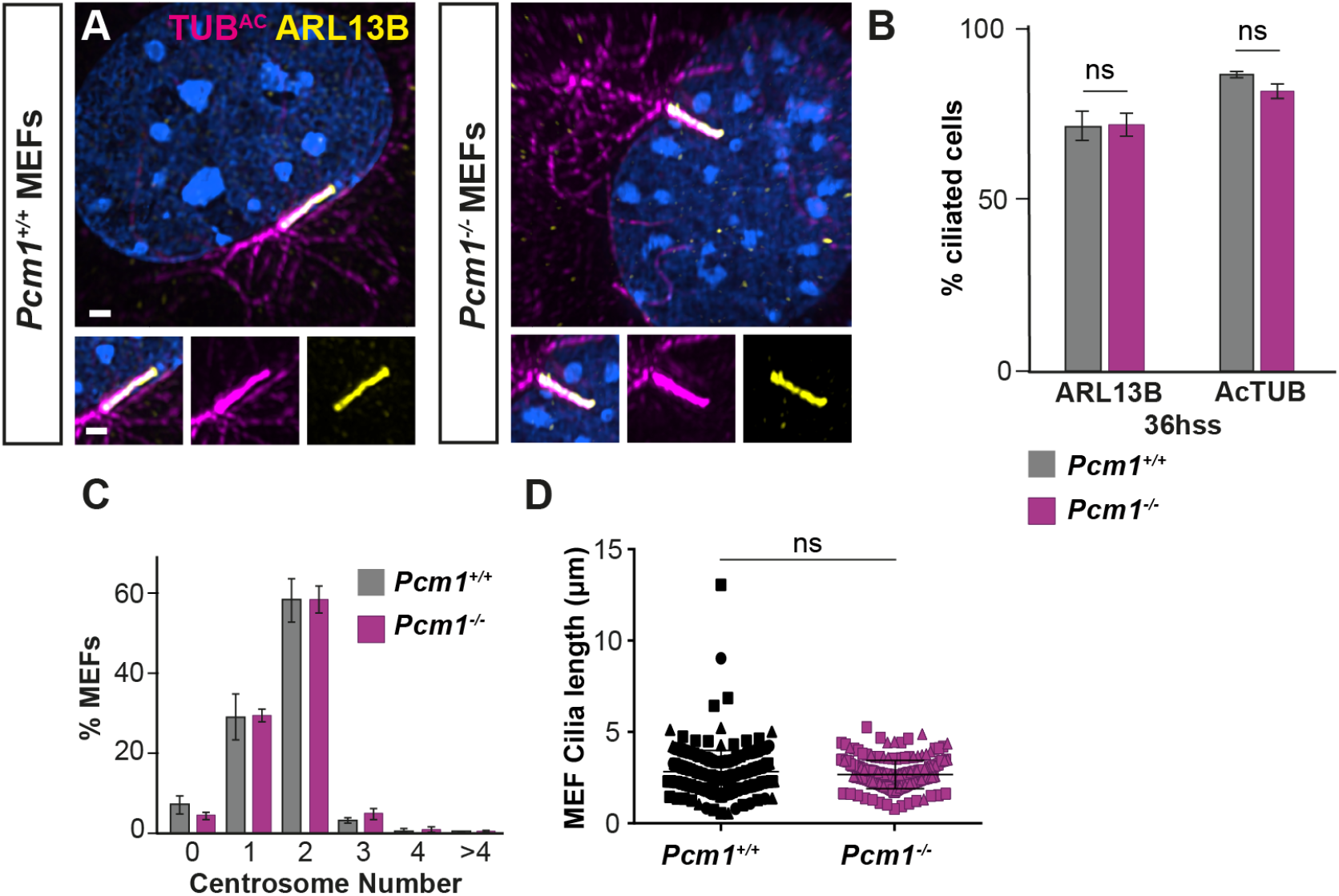
PCM1 is dispensable for ciliogenesis in MEFs. **(A)** Wild-type and *Pcm1*^−/−^ MEFs serum starved for 36 h and immunostained for ARL13B (yellow), TUB^*Ac*^ (magenta) and nuclei (DAPI, blue). Scale bar represents 10 *μ*m. **(B)** Quantification of ciliogenesis in wild-type and *Pcm1*^−/−^ MEFs serum immunostained as in A. Error bars represent SEM, n=3 MEF lines. Student’s t-test ns: not significant. Ciliogenesis is unaffected in *Pcm1*^−/−^ MEFs. **(C)** Quantification of centrosome number in wild-type and *Pcm1*^−/−^ MEFs serum immunostained as in Figure 4I. Error bars represent SEM, n=3 MEF lines. One way ANOVA. Centrosome duplication is unaffected in *Pcm1*^−/−^ MEFs. **(D)** Quantification of ciliary length in wild-type and *Pcm1*^−/−^ MEFs serum immunostained for ARL13B. Student’s t-test ns: not significant. PCM1 is not critical for cilia length regulation.

**Figure 5–Figure supplement 1.**
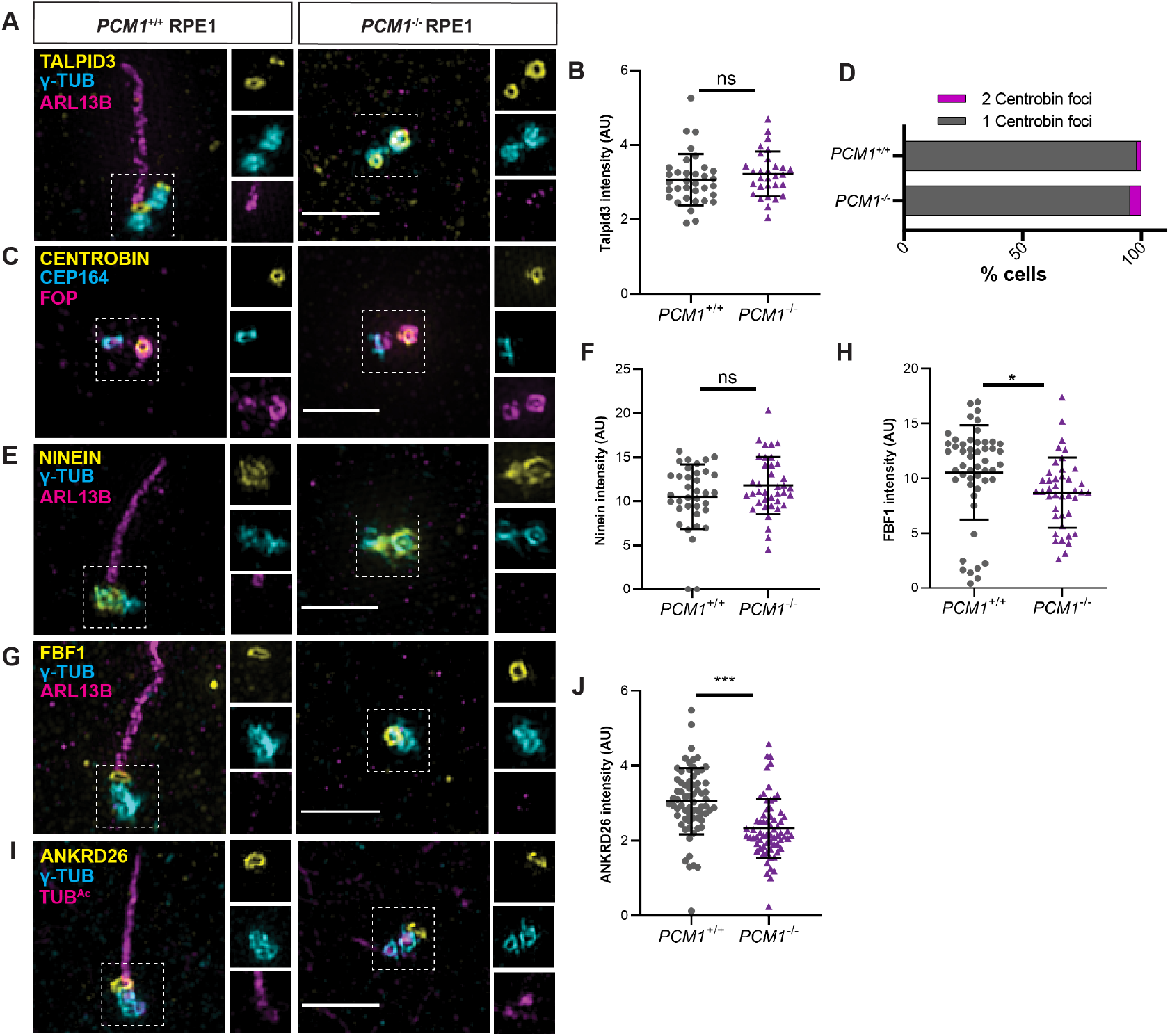
PCM1 is dispensable for mother centriole maturation. (A) Wildtype and *PCM1*^−/−^ RPE1 cells immunostained for TALPID3 (yellow), centrioles (*γ*-tubulin, cyan), and cilia (ARL13B, magenta). (B) Quantification of TALPID3 levels at centrioles. PCM1 is not required for TALPID3 localization to the distal centriole of RPE1 cells. (C) Immunostaining for Centrobin (yellow), distal appendages (CEP164, cyan) and centrioles (FOP, magenta). (D) Quantification of number of Centrobin foci at centrioles per cell. PCM1 is dispensable for Centrobin localization to the daughter centriole, identified by the absence of distal appendages. (E) Immunostaining for subdistal appendage protein Ninein (yellow), centrioles (*γ*-tubulin, cyan), and cilia (ARL13B, magenta). (F) Quantification of Ninein levels at centrioles. PCM1 is not required for Ninein localization to subdistal appendages. (G) Immunostaining for distal appendage protein FBF1 (yellow), centrioles (-tubulin, cyan), and cilia (ARL13B, magenta). (H) Quantification of FBF1 levels at mother centrioles. FBF1 is modestly reduced at the distal appendages of *PCM1*^−/−^ RPE1 cells. (I) Immunostaining for distal appendage protein ANKRD26 (yellow), centrioles (-tubulin, cyan), and cilia (TUB^*Ac*^, magenta). (J) Quantification of ANKRD26 levels at mother centrioles. ANKRD26 is also modestly reduced at the distal appendages of *PCM1*^−/−^ RPE1 cells. Scale bars represent 2 *μ*m, and in insets represent 0.5 *μ*m. Student’s t-test, * P<0.05, *** P<0.001, ns: not significant.

**Figure 5–Figure supplement 2.**
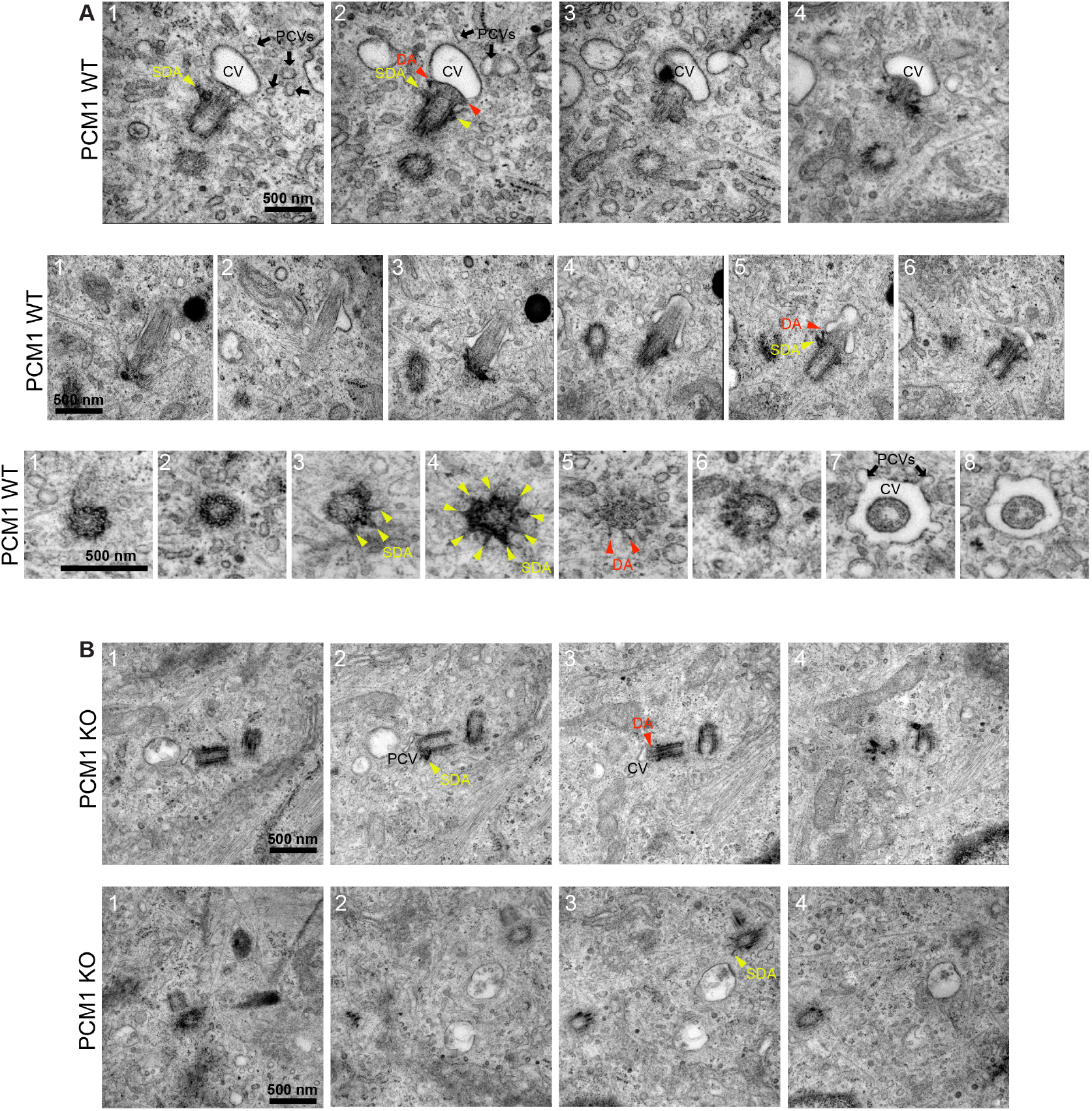
PCM1 promotes mother centriole association with vesicles. Serial-section TEM images of wild-type (A) and *PCM1*^−/−^ (B) RPE1 cells 1 h post serum starvation. In both wild-type and *PCM1*^−/−^ cells, mother centrioles possess subdistal (SDA, yellow arrowheads) and distal appendages (DA, red arrowheads). (A) In wild-type cells, the mother centriole is associated with preciliary vesicles (PCVs, black arrows) and ciliary vesicles (CV). (B) In *PCM1*^−/−^ RPE1 cells, the mother centriole is associated with few PCVs or CVs. Scale bars represent 500 nm.

**Figure 7–Figure supplement 1.**
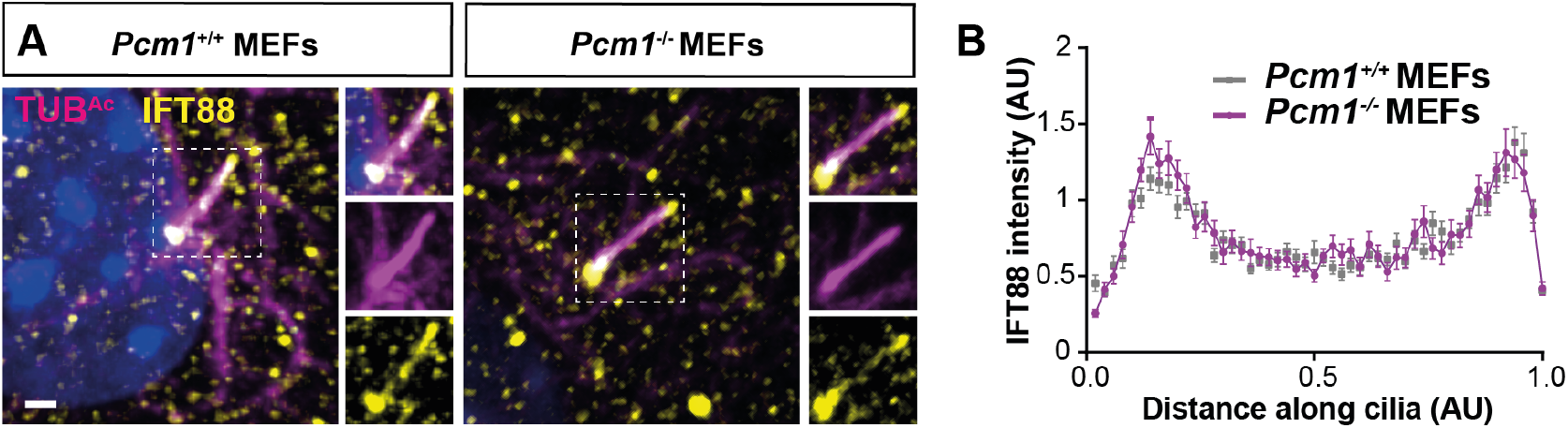
PCM1 does not control IFT88 levels in MEF cilia. **(A)** Wild-type and *Pcm1*^−/−^ MEFs immunostained for IFT88 (yellow), cilia (TUB^*Ac*^, magenta) and nuclei (DAPI, blue). **(B)** Quantification of IFT88 intensity along the cilia, calculated from line plots of >120 wild-type and *Pcm1*^−/−^ cilia from 3 primary cell lines per genotype. IFT88 levels in cilia is not highly dependent on PCM1.

**Figure 8–Figure supplement 1.**
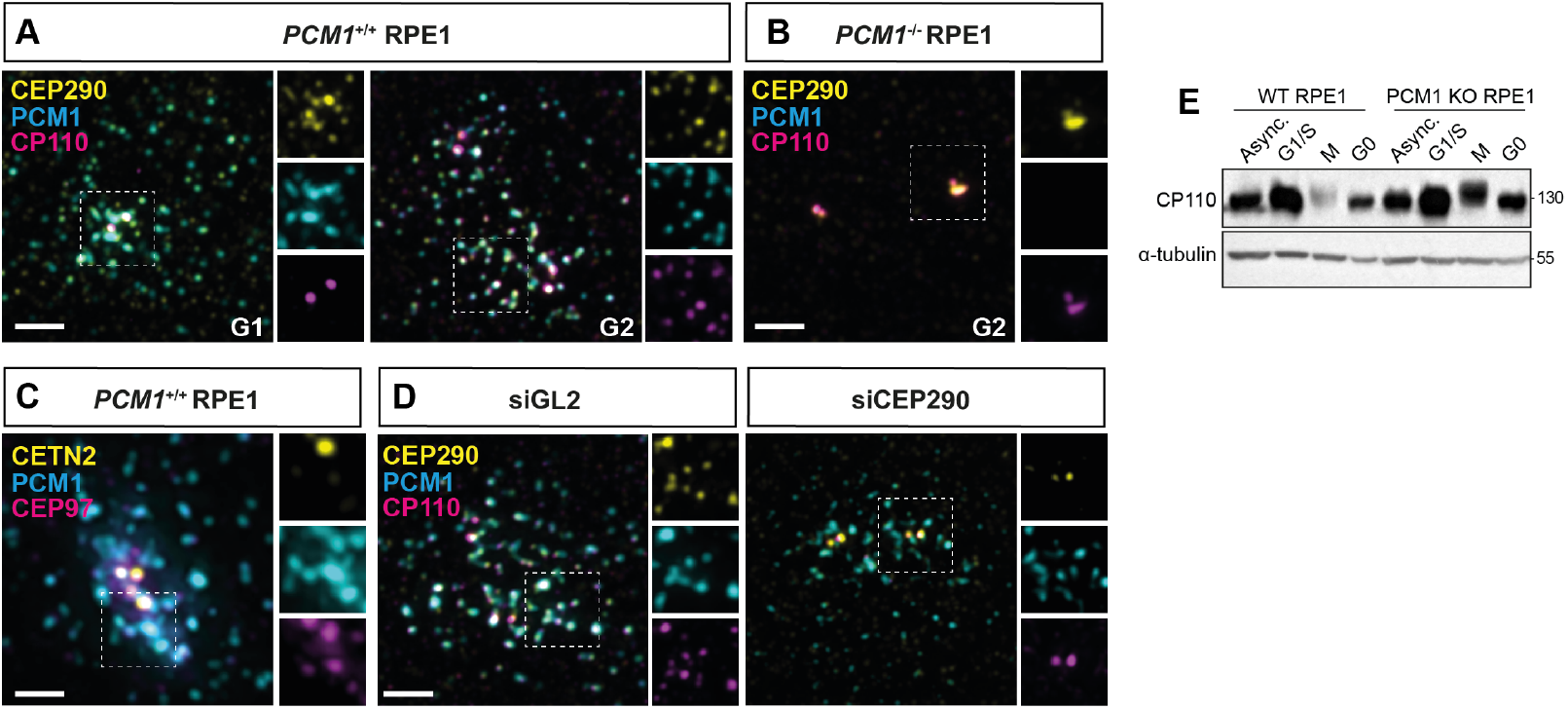
CP110 localizes to satellites in a CEP290-dependent manner. **(A, B)** CP110 localizes to satellites in cycling RPE cells at G1 and G2, and this satellite pool is lost in *PCM1*^−/−^ RPE cells (B). **(C)** Satellite localization of CP110 is dependent on CEP290; siRNA knockdown of *CEP290* leads to loss of the CP110 satellite pool. **(D)** Western blot demonstrating that cell cycle dependent degradation of CP110 in mitosis is disrupted in *PCM1*^−/−^ RPE1 cells. Scale bars represent 2 *μ*m.

